# Interpretable AI-driven materiomics to decode microenvironmental cues for stem cell immunomodulation

**DOI:** 10.64898/2026.02.26.708394

**Authors:** Chenxi Pan, Yi He, Yingying Duan, Kaiwen Chen, Qifan Wang, Xiangying Wang, Yonggang Zhang, Chuanfeng An, Huanan Wang

## Abstract

Hydrogels that mimic the extracellular matrix create microenvironments containing diverse physicochemical cues that regulate stem cell fate, particularly their immunomodulatory and pro-regenerative functions. However, elucidating how microenvironmental cues regulate cell fate remains challenging because of complex multi-parameter interactions and nonlinear relationships, thereby requiring large numbers of samples and substantial experimental effort. Here, we present a materiomics platform that integrates a large-scale systematic dataset with artificial intelligence (AI) to evaluate and predict optimal stem cell niche features for enhancing the immunomodulatory functions of mesenchymal stem cells (MSCs). Specifically, binary hydrogels composed of methacrylated alginate and gelatin (AMGM) were used to construct a comprehensive materiomics database incorporating seven critical physicochemical cues associated with MSC-mediated regulation of macrophage polarization. Machine learning models trained on this materiomics database successfully predicted hydrogel formulations that enhanced MSC immunomodulatory efficacy and further identified matrix stiffness as the physicochemical cue with the greatest contribution to MSC-mediated immunomodulation within the AMGM hydrogel system. In summary, this study demonstrates the utility of AI for evaluating the relative contributions of microenvironmental cues and establishes a data-driven research framework for investigating microenvironmental regulation of stem cell function.

## 1. Introduction

Mesenchymal stem cells (MSCs) possess unique immunomodulatory and pro-regenerative capacities, making them a promising therapeutic strategy for treating inflammatory conditions in regenerative medicine [1]. Their ability to regulate immune responses, mainly through paracrine signaling, is now recognized as a major contributor to their therapeutic efficacy in numerous regenerative conditions [1, 2]. The unique immunomodulatory properties of MSCs have been previously demonstrated to mediate the proliferation, apoptosis, migration, and function of several immune cell types, such as macrophages, neutrophils, natural killer cells, T cells, and B cells [3]. Among these immune cells, macrophages are central effectors of the innate immune system and play critical roles in inflammation and tissue repair through dynamic transitions between pro-inflammatory (M1) and anti-inflammatory (M2) phenotypes in response to microenvironmental cues [4]. MSCs have been shown to promote tissue healing by attracting monocytes and macrophages and inducing macrophages to polarize towards the M2 phenotype [1], which also has a profound impact on the host inflammatory responses upon the implantation of external biomaterials [5]. Consequently, engineering the immune microenvironment to influence both stem cell and macrophage behavior has become a significant focus of regenerative medicine research.

Hydrogels serving as biomimetic extracellular matrices (ECMs) can establish an engineered microenvironment for the culture and regulation of MSC functionalities, including preserving cell stemness and secretory capacity, improving cell survival and local retention, achieving targeted delivery efficiency, and allowing precise control of secretory profiles [6]. Due to the high degree of control over the physicochemical cues of hydrogels, they have been widely used as a platform for functional and mechanistic investigations of MSC behavior. For instance, matrix stiffness has recently been recognized as a key regulatory factor influencing the secretory functions of stem cells and, consequently, their therapeutic efficacy. Our prior study revealed the impact of the stiffness of gelatin methacryloyl (GelMA) hydrogels on the immunomodulatory capacity of encapsulated MSCs, as reflected by macrophage polarization and associated inflammatory responses [7]. Although these studies have demonstrated that matrix stiffness and viscoelasticity influence stem cell paracrine signaling and immune regulation [6–8], the roles of RGD ligand spacing and surface charge, which are primarily studied in stem cell differentiation, remain less understood in paracrine immunomodulation [8–11].

Specifically within differentiation models, recent pioneering research has revealed that extended RGD nanospacing exceeding 70 nm paradoxically promotes osteogenesis despite disrupting integrin clusters and impairing initial cell adhesion. This paradigm-shifting phenomenon occurs because weak adhesion promotes the nuclear translocation and polymerization of globular actin, increasing nuclear tension and thereby activating osteogenic genes expression independently of conventional myosin-based traction forces [11]. In immunomodulatory contexts, however, matrix effects are even more difficult to interpret because synthetic niches comprise multiple coupled physicochemical cues rather than stiffness alone [12, 13]. Moreover, these cues often influence immune outcomes in a non-monotonic manner. For example, hydrogels with intermediate stress-relaxation times promoted more pronounced M2 macrophage polarization than either fast- or slow-relaxing matrices [14], whereas precision-templated scaffolds with pore sizes of approximately 30-40 μm elicited more favorable healing-associated responses than scaffolds with either smaller or larger pores [15]. Together, these observations underscore the multidimensional and nonlinear nature of matrix regulation, complicating efforts to resolve the specific contribution of individual matrix properties to stem cell paracrine immunomodulation. Because multiple microenvironmental cues vary simultaneously and often interact nonlinearly, identifying optimal hydrogel formulations requires extensive experimental screening and remains highly challenging. These challenges highlight the need for systematic and efficient strategies capable of defining experimentally relevant design spaces while resolving multifactorial regulatory mechanisms.

In recent years, materiomics has emerged as an interdisciplinary field that integrates materials science, genomics, data science, and artificial intelligence to accelerate biomaterial discovery [16]. Its central goal is to decode the relationships among material composition, structure, properties, and function through high-throughput experimentation, multidimensional data integration, and machine learning, thereby shifting biomaterial design from empirical trial-and-error approaches toward data-driven optimization [17, 18]. Existing AI-assisted strategies in biomaterial research can be broadly divided into two categories. One category uses high-dimensional biological data, such as transcriptomic profiles, to predict cell fate or differentiation outcomes. For example, the Deng group employed 12 transcriptomic datasets to predict MSC differentiation with 90% accuracy using a k-nearest neighbor classifier, enabling outcome prediction as early as day 7 [19]. Similarly, the Wang group used RNA-sequencing data collected from MSCs cultured in hydrogels to model myogenic and osteogenic differentiation, although the predictive performance of their random forest and linear regression models remained limited [20]. While these studies are valuable for early biological prediction, they mainly establish correlations between omics features and cell fate, and provide limited insight into how intrinsic biomaterial properties regulate the target function. The second category uses material composition as model inputs to predict intrinsic material performance. For example, the Chen group used material composition to predict photothermal conversion efficiency with an XGBoost-assisted model [21]. Although effective for optimizing material centered properties, such studies do not address how intrinsic biomaterial properties regulate biologically relevant cell functions. Therefore, there remains a need for machine-learning studies that directly link the intrinsic physicochemical properties of biomaterials to biologically relevant MSC function.

To address this gap, the present study established an integrated materiomics platform that directly links hydrogel composition, matrix-related physicochemical cues, and MSC immunomodulatory function within one unified framework. Binary hydrogels composed of methacrylated alginate and methacrylated gelatin (AMGM) were fabricated via free-radical photopolymerization, generating a library containing more than one hundred formulations. This material system enabled systematic variation of seven key physicochemical cues, including matrix stiffness, RGD spacing, surface charge, stress relaxation, pore size, hydrophilicity/hydrophobicity, and viscosity. By combining this multiparameter hydrogel database with machine learning analysis, we sought not only to predict hydrogel formulations that enhance MSC-mediated immunomodulatory efficacy, but also to compare the relative contributions of multiple coupled matrix cues. In this way, the present work provides a material-property-function framework for investigating how intrinsic hydrogel properties regulate stem cell immunomodulation, while also reducing the experimental burden associated with conventional optimization strategies that vary one factor at a time (Figure 1).

**Figure 1.**
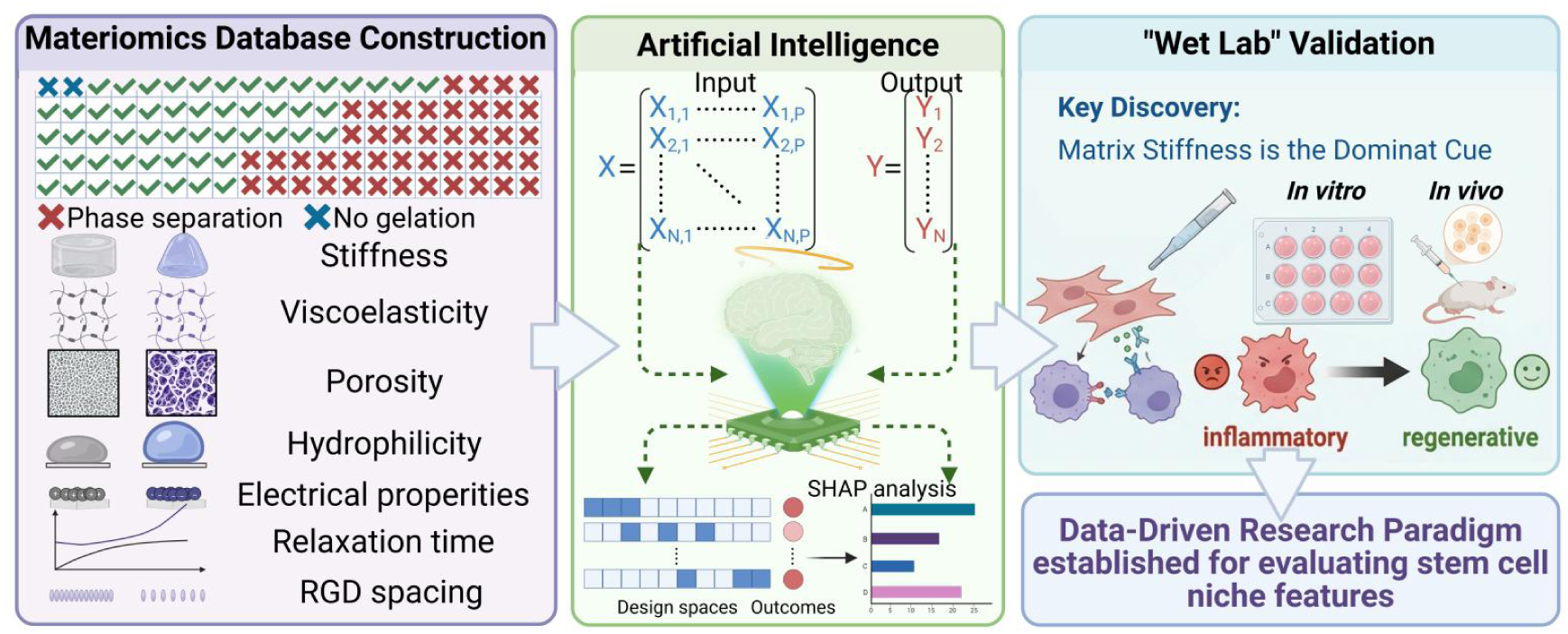
Workflow illustrating the machine learning based study design for investigating and validating the effects of physicochemical cues on stem cell immunomodulation.

## 2. Results

### 2.1 Establishment of a Materiomics Database Based on AMGM Hydrogel Matrices with Varying Compositions

To establish a biocompatible and mechanically tunable microenvironment for mesenchymal stem cell immunomodulation, we selected a hybrid hydrogel system based on methacrylated alginate (AlgMA) and methacrylated gelatin (GelMA). Both alginate and gelatin are naturally derived polymers that have been extensively used in tissue engineering and cell microenvironment studies [22, 23]. GelMA provides ECM mimetic features, including intrinsic cell-adhesive motifs and matrix metalloproteinase-responsive characteristics, whereas AlgMA offers a readily processable and compositionally versatile polymer framework [22, 23]. After methacrylation, both components can be efficiently crosslinked to form hydrogel networks, thereby enabling systematic tuning of matrix stiffness and viscoelasticity. In addition, alginate-containing networks broaden the accessible mechanical design space, while the combination of AlgMA and GelMA enables the generation of relatively homogeneous hybrid matrices across a wide compositional range, making this system well suited for combinatorial matrix design.

The precursor concentration ranges were established through preliminary screening and experimental optimization. Specifically, we selected 0.5-2.5% w/v for AlgMA and 0.5-10% w/v for GelMA because these ranges consistently supported hydrogel network formation, provided a broad yet experimentally manageable spectrum of physicochemical properties, and maintained acceptable cellular viability during extended culture (Supplementary Figure 2). These concentration ranges were therefore chosen to balance matrix tunability, structural integrity, and cytocompatibility.

To construct a photo-crosslinkable binary hydrogel system, we first synthesized AlgMA and GelMA by chemically modifying sodium alginate and gelatin with methacrylic anhydride. The two components were then crosslinked via free-radical photopolymerization under 365 nm UV irradiation (250 mW/cm², 60 s) using 0.2% w/v LAP as the photoinitiator to form AMGM hydrogels (Figure 2A). Fourier transform infrared spectroscopy (FT-IR) confirmed the successful synthesis of AlgMA and GelMA, showing characteristic absorption peaks at 1730 cm^-1^ and 1633 cm^-1^, respectively (Supplementary Figure 1A, B). The degrees of methacrylation (DOM) were determined by ¹H NMR using the sugar-ring H^-1^ peaks for AlgMA and the lysine ε-methylene H^-1^ peaks for GelMA used as internal standards. The calculated DOM values were 47.7% for AlgMA and 85.3% for GelMA (Supplementary Figure 1C, D). To prepare AMGM hydrogel matrices with varied compositions, precursor solutions containing 0.5-2.5% w/v AlgMA and 0.5-8% w/v GelMA were mixed and polymerized upon UV initiation. Three distinct states were observed: (i) phase separation, (ii) non-gelation, and (iii) formation of uniform binary hydrogels (Figure 2B, C; Supplementary Figure 2). Among the AM*x*GM*y* binary combinations, 54 formulations consistently yielded uniform hydrogels, thereby forming the basis of the materiomics database used in subsequent analyses (Figure 2D).

**Figure 2.**
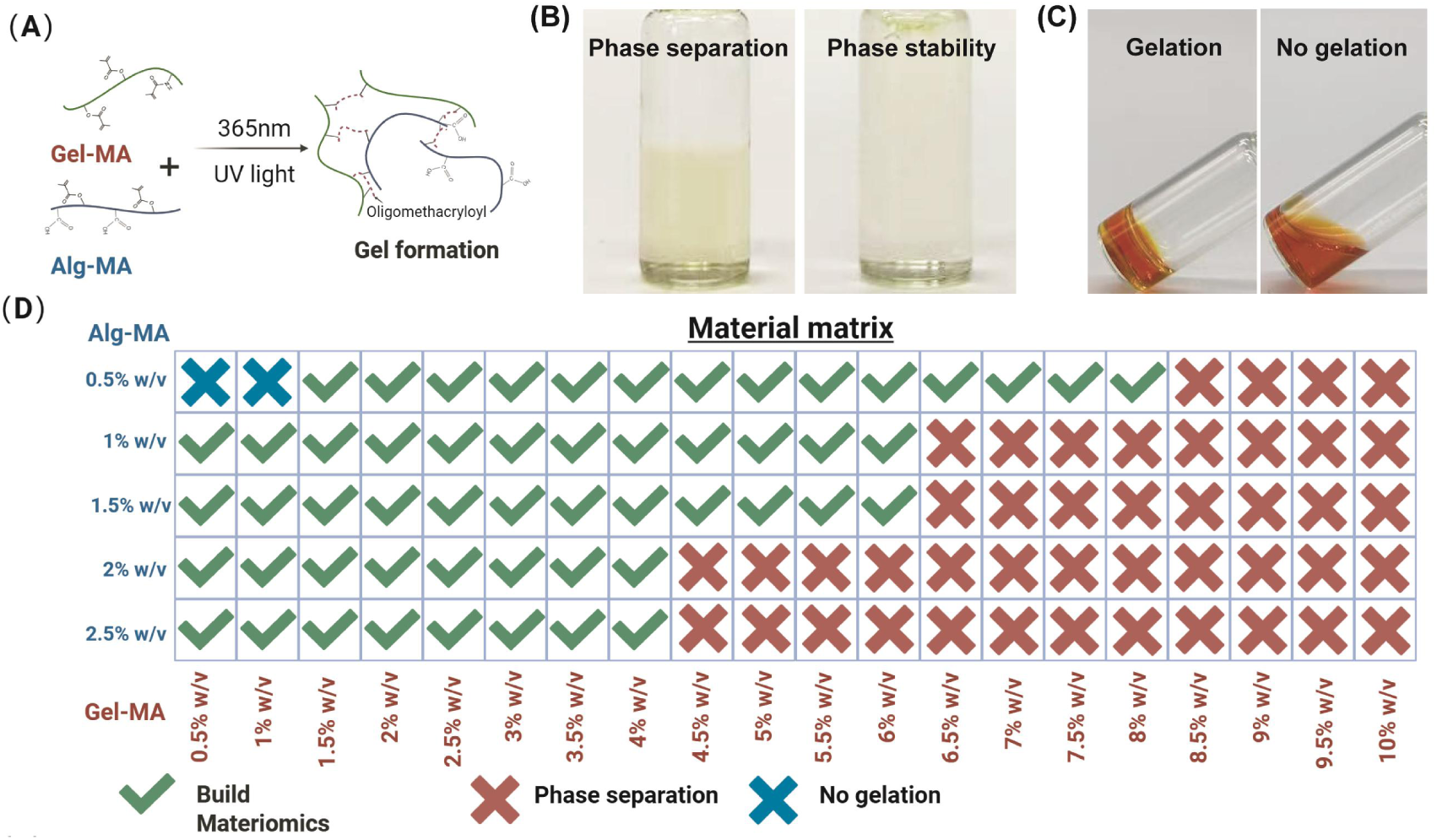
Construction of the AMGM Hydrogel database. (A) Schematic illustration of the synthesis of GelMA (gelatin methacryloyl) and AlgMA (alginate methacryloyl). (B) Representative image of phase separation observed in hydrogel precursor solutions. (C) Representative image of non-gelation observed in hydrogel precursor solutions. (D) Screening of AMGM formulations. Valid hydrogel compositions were identified by excluding formulations exhibiting phase separation or non-gelation.

Rational design of the compositional and structural properties of biomaterials serving as ECM enables precise regulation of biophysical and biochemical microenvironmental cues, thereby modulating MSC behavior and functions, including secretory activity [6]. From surface charge, chemical functional groups, and adhesive ligand presentation at the molecular level, to surface topography and pore size at the microstructural level, and ultimately to bulk mechanical properties at the macrostructural level, all these factors have been verified to influence MSC fate [22]. In the present binary AMGM hydrogel system, seven key variables, including surface charge, RGD spacing, hydrophilicity/hydrophobicity, pore size, matrix stiffness, viscoelasticity, and stress relaxation time, can be systematically tuned and are expected to exert significant effects on MSC function.

As an important surface characteristic, biomaterial surface charges can directly impact cell-material interactions, including cell adhesion and migration, protein adsorption, host foreign-body responses, and cell-mediated material internalization and degradation [10]. In the present study, AMGM hydrogels exhibited zeta potentials ξ ranging from -5.54 mV to 4.98 mV (Figure 3A). We further hypothesized that this relatively low charge may correlate with the high grafting rate of AlgMA and GelMA, as this grafting efficiency could diminish the density of charged groups on the material surface.

**Figure 3.**
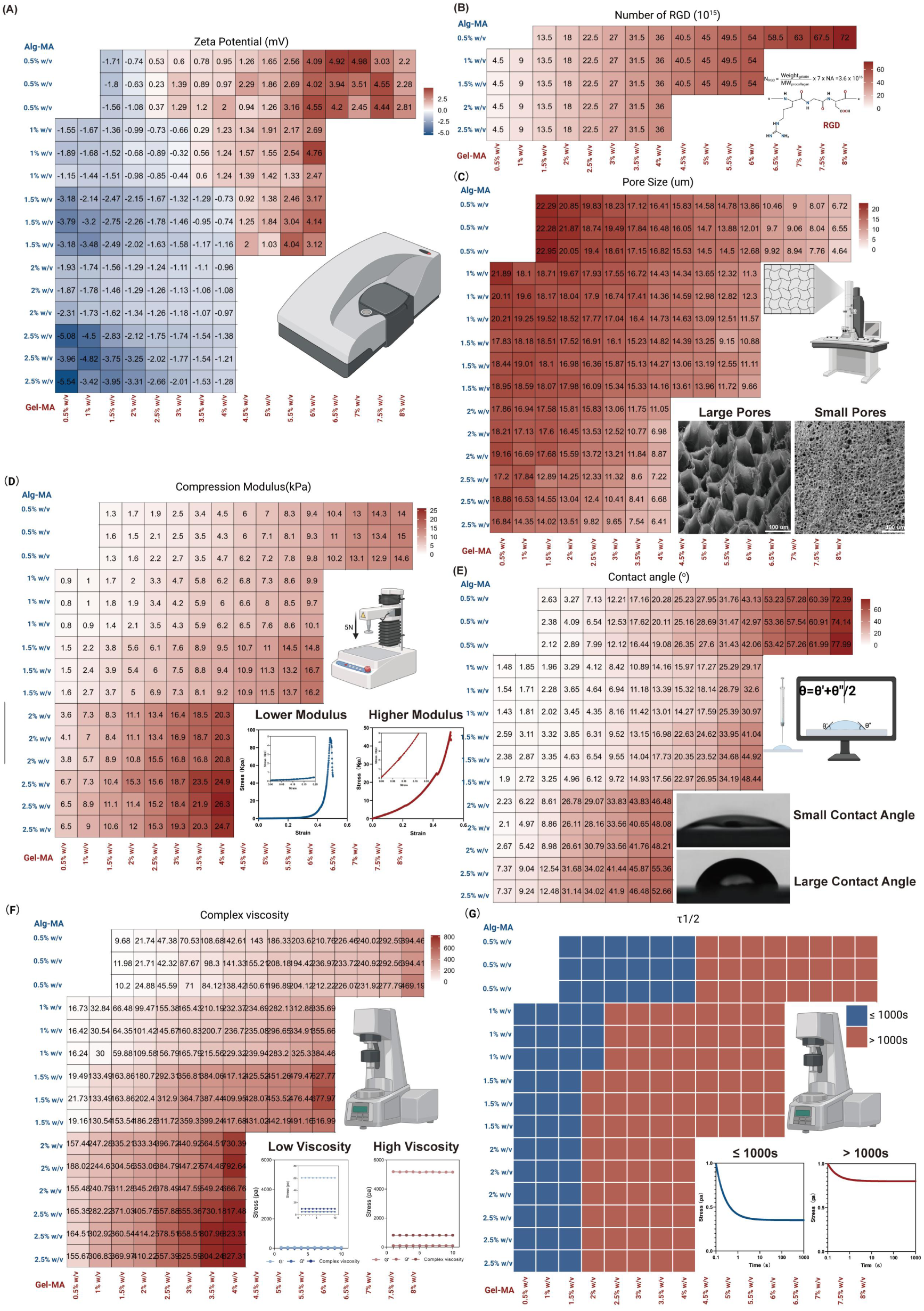
Physicochemical characterization of the AMGM hydrogel database. (A) Charge density characterization, (B) RGD sequence density determination, (C) pore size characterization, (D) surface contact angle measurement, (E) Young’s modulus measurement, (F) complex viscosity quantitative analysis, and (G) stress relaxation half-relaxation time (τ₁/₂) of AMGM hydrogels with different compositions.

The presence of RGD sequences is crucial for cell adhesion, migration, and differentiation [9]. To investigate the role of RGD sequences in MSC immunomodulatory function, we utilized binary AMGM hydrogel systems in which GelMA intrinsically provides RGD sequences into the binary AMGM hydrogels, achieving systematic modulation of RGD spacing and density by varying polymer concentrations [23]. Consequently, we found that the RGD sequence density within the AMGM hydrogel system ranged from 4.5×10¹⁵ to 7.2×10¹⁶ molecules/cm² (Figure 3B). As expected, RGD sequences density increased with GelMA content.

Pore size is an important structural feature of biomaterials that influences cellular and tissue responses in tissue engineering. Previous studies have demonstrated that different pore sizes can support distinct tissue regeneration requirements [24]. Specifically, pore sizes of around 5 μm promote neovascularization, those of 20-125 μm facilitate skin regeneration, and those of 100-350 μm support bone regeneration [25]. In this study, the pore sizes of AMGM hydrogels were evaluated by scanning electron microscopy (SEM) imaging of lyophilized hydrogels. The analysis revealed pore sizes ranging from 4.64 to 22.95 μm. Moreover, pore size was negatively correlated with total polymer content. Specifically, formulations with lower polymer content exhibited larger pores, whereas increasing polymer content resulted in denser network structures with smaller pore sizes (Figure 3C).

As another fundamental property of biomaterials, intrinsic hydrophilicity/hydrophobicity significantly influence cellular behavior. Current research exploring the effects of hydrophilic/hydrophobic materials with MSC biology have focused primarily on the maintenance of stemness. Specifically, recent advancements demonstrate that three dimensional (3D) culture of hematopoietic stem and progenitor cells within highly hydrophilic zwitterionic hydrogels substantially maintains their cellular stemness [26]. However, the influence of material hydrophilic/hydrophobic on the immunomodulatory functions of MSCs remains largely unexplored.

To address the research gap resulting from the limited number of studies investigating the relationship between material hydrophilicity/hydrophobicity and MSC immunomodulation, we performed contact angle measurements and found that the binary hydrogel matrices exhibited substantial variations in hydrophilicity, with contact angles ranging from 1.48° to 77.99°. These wide variations reflect the diverse hydrophilic and hydrophobic characteristics of hydrogels with different compositions (Figure 3E).

Over the past decade, matrix stiffness has been widely recognized as a key mechanical parameter governing MSC fate by resisting cell-generated traction forces. However, many of these studies implicitly assume that the native MSC niche behaves as a purely elastic material. In reality, natural ECM, as well as most tissues and organs, exhibit viscoelastic behavior across multiple time scales [27, 28]. Unlike purely elastic materials, viscoelastic matrices dissipate applied forces, thereby altering how cell-generated traction forces are transmitted, stored, and remodeled within the microenvironment, as well as how external mechanical signals are conveyed to intracellular mechanotransduction pathways [12]. Therefore, investigations of MSC mechanobiology should consider both the elastic and viscoelastic properties of synthetic niches.

Previous studies in MSC mechanobiology have identified matrix stiffness as a key regulator of MSC fate, with soft matrices favoring neurogenesis, matrices of intermediate stiffness promoting myogenesis, and rigid matrices enhancing osteogenesis [29]. In addition, matrix stiffness has been shown to influence MSC immunomodulatory function, macrophage polarization, and the associated inflammatory responses [7, 30, 31]. To characterize the mechanical properties of AMGM hydrogels, we measured their Young’s modulus via compression tests, and the measured values ranged from 0.8 to 26.3 kPa, successfully spanning the stiffness ranges of diverse tissue microenvironments (Figure 3D).

Native ECM does not behave as a perfectly elastic solid. Instead, when subjected to mechanical deformation, reconstituted ECM initially resists deformation and subsequently undergoes time-dependent energy dissipation, a hallmark of viscoelastic materials [32]. This energy dissipation arises from the dynamic molecular organization of the ECM, which is not composed of a perfectly crosslinked network but instead undergoes weak bond dissociation, protein unfolding, and entanglement release [33–35]. Therefore, viscoelasticity is also a significant indicator of MSC fate. Therefore, viscoelasticity represents another important matrix property influencing MSC fate. To characterize matrix viscoelasticity in AMGM hydrogels, we measured complex viscosity and found values ranging from 9.68 to 827.31 Pa·s, indicating substantial differences in viscoelastic behavior among the formulations (Figure 3F). In addition, stress relaxation was evaluated using the half-relaxation time (τ₁/₂), another key descriptor of matrix viscoelasticity [36]. Using 1000 s as a threshold, more than 60% of the formulations exhibited τ₁/₂ values above this value and were therefore classified as slow-relaxing matrices, whereas the remaining formulations displayed relatively rapid stress-relaxation behavior (Figure 3G).

Collectively, these data enabled the establishment of a materiomics database comprising 54 binary AMGM formulations, 162 samples, and 1134 data points describing seven physicochemical parameters associated with each formulation. Systematic characterization of this material library revealed how these physicochemical properties varied across the compositional space. By restricting the composition layer to formulations that satisfied the requirements for cell-laden hydrogel culture, we were able to generate a physicochemical dataset within a standardized and experimentally relevant range. This strategy provided a sufficiently diverse dataset for subsequent machine-learning analysis while maintaining a manageable experimental workload.

### 2.2 Quantitative Evaluation of MSC-Mediated Macrophage Polarization Using the AMGM Hydrogel Database

To evaluate the suitability of AMGM hydrogels as functional 3D niches for MSC delivery, we investigated the biological behavior of living cells encapsulated within these matrices. Specifically, umbilical cord-derived mesenchymal stem cells (UCMSCs) were homogeneously suspended in precursor solutions and subsequently encapsulated through photopolymerization under UV irradiation (365 nm, 250 mW/cm², 60 seconds) (Figure 4A). To systematically evaluate the cytocompatibility of the encapsulation process, live/dead fluorescent staining was performed. The results showed that UCMSCs maintained high viability across all selected hydrogel formulations after 48 h of culture (Figure 4C). To further evaluate cell proliferation, a CCK-8 assay was performed. The optical density values increased continuously over time, showing an approximately threefold increase between 24 and 72 h (Figure 4B), suggesting progressive cell growth within the hydrogel matrices. Together, these results indicate that the AMGM hydrogels provide a cytocompatible microenvironment that supports MSC survival and proliferation under the tested conditions.

**Figure 4.**
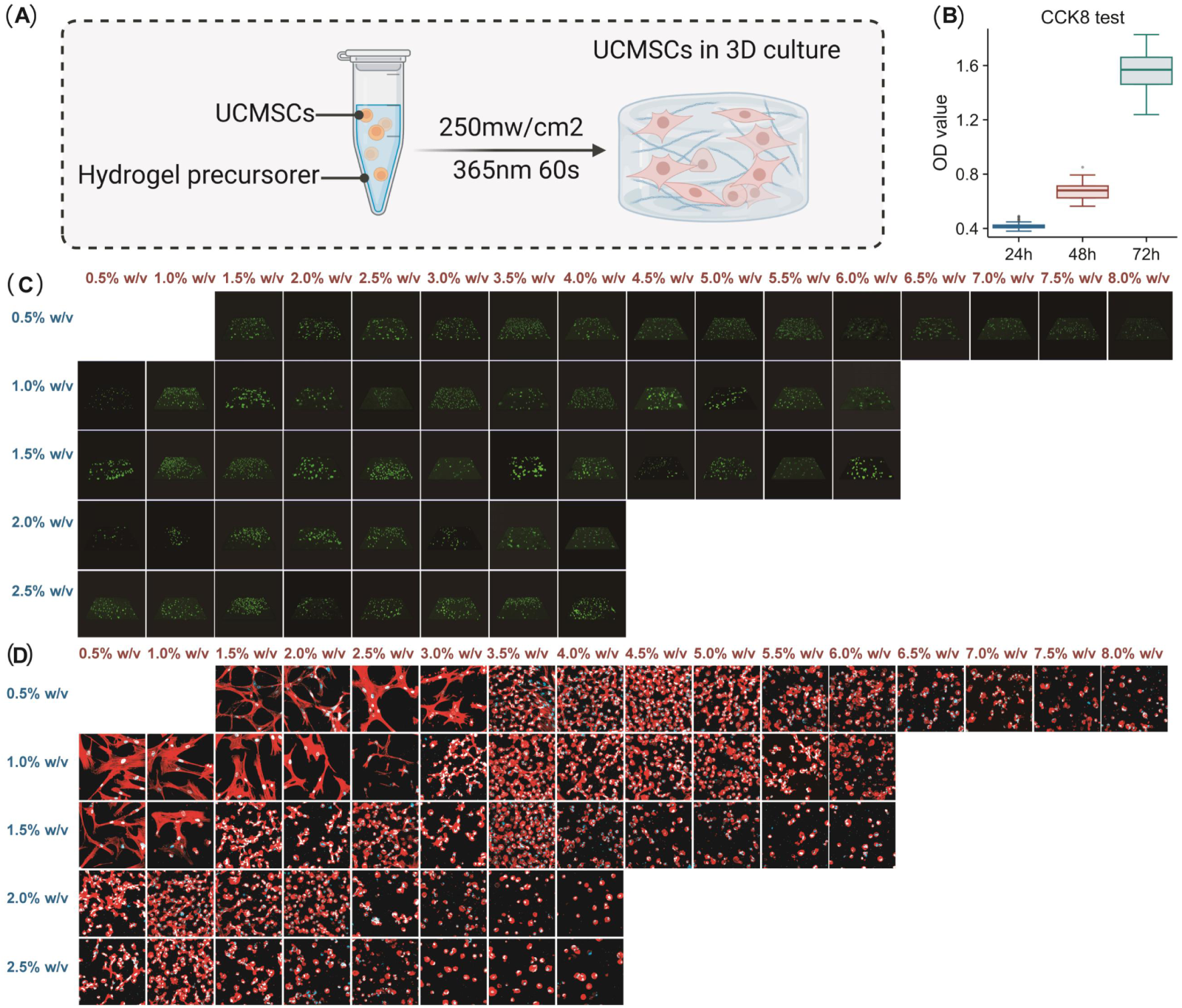
UCMSC responses within the AMGM hydrogel database. (A) Schematic illustration of UCMSC encapsulation within three-dimensional AMGM hydrogels. (B) Cell viability analysis of UCMSCs cultured in different hydrogel formulations for 24, 48, and 72 h (n = 162). (C) Representative live/dead fluorescence images of UCMSCs cultured in different hydrogel formulations for 48 h (n = 54). (D) Cytoskeletal staining and morphological characterization of UCMSCs cultured in different hydrogel formulations (n = 54).

Beyond supporting cell viability, the engineered ECM also provides structural cues that regulate cell morphology and behavior. To evaluate the ability of UCMSCs to respond to the surrounding microenvironment, cytoskeletal staining was performed on cells encapsulated within different AMGM hydrogel formulations (Figure 4D). Through morphological analysis, we observed substantial differences in the cytoskeletal organization of UCMSCs encapsulated within different hydrogel formulations [7]. MSCs cultured in hydrogels with lower polymer content exhibited a more spread morphology with extensive cellular protrusions, whereas those encapsulated in higher-polymer-content hydrogels appeared more rounded. Quantitative analysis of the cytoskeletal aspect ratio confirmed a progressive reduction in cell elongation with increasing polymer content (Supplementary Figure 3). These observations demonstrate that variations in hydrogel composition and matrix properties regulate MSC morphology within the 3D microenvironment.

Phenotypic characterization confirmed that the isolated UCMSCs exhibited the expected MSC phenotype, consistent with previous reports [37]. Specifically, the cells expressed the mesenchymal markers CD105, CD73, and CD90, while showing negligible expression of the hematopoietic marker CD34 (Figure 5A). Macrophage phenotypes were also validated prior to co-culture experiments. Naïve macrophages (M0) expressed the pan-macrophage marker F4/80. Following polarization, classically activated macrophages (M1) expressed the pro-inflammatory markers CD80 and CD86, whereas alternatively activated macrophages (M2) expressed the anti-inflammatory marker CD206 (Figure 5B). These results confirmed the successful establishment of the macrophage polarization model used to evaluate MSC-mediated immunomodulatory function.

**Figure 5.**
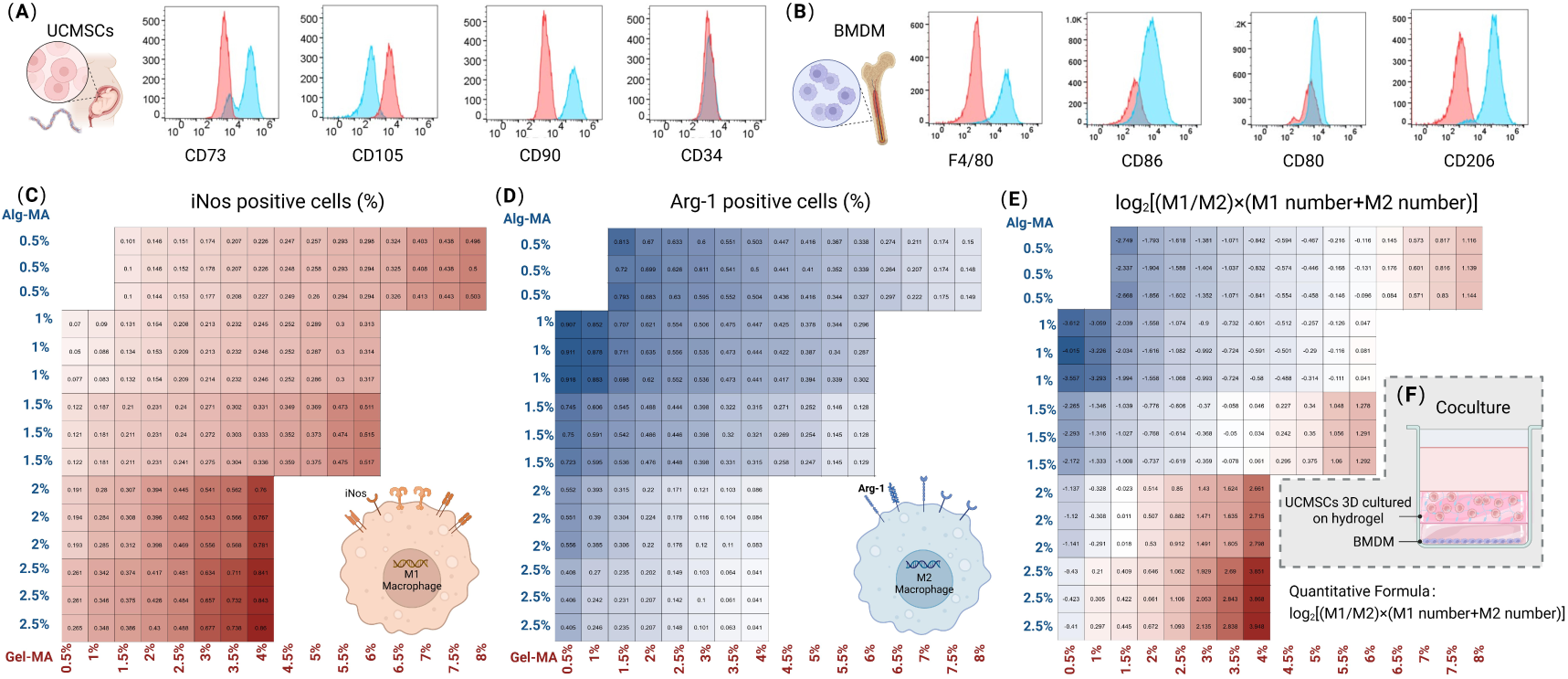
Evaluation of MSC-mediated macrophage polarization. (A) Phenotypic characterization of umbilical cord-derived mesenchymal stem cells (UCMSCs) using CD73, CD105, CD90, and CD34 markers. (B) Phenotypic characterization of bone marrow-derived macrophages using F4/80, CD80, CD86, and CD206 markers. (C-E) Results of indirect co-culture between cell-laden hydrogels and macrophages. (C) Quantification of iNOS-positive macrophages (n = 162). (D) Quantification of Arg-1-positive macrophages (n = 162). (E) Distribution of the macrophage polarization index calculated using the standardized quantitative formula (n = 162). (F) Schematic illustration of the Transwell co-culture system used to assess MSC-mediated macrophage polarization, including the quantitative index applied for data analysis.

Macrophages are the primary cell type involved in the immune response and can guide regenerative processes by sensing environmental stimuli and polarizing into distinct phenotypic states [38]. To further investigate how the UCMSCs encapsulated within different AMGM hydrogel formulations regulate macrophage polarization, a Transwell co-culture system was established in which 3D-encapsulated UCMSCs were indirectly cultured with macrophages (Figure 5F). To better mimic inflammatory conditions relevant to tissue repair, M0 macrophages were first polarized toward an M1 phenotype before co-culture. Macrophage phenotypic responses were subsequently evaluated by immunofluorescence staining of the pro-inflammatory marker inducible nitric oxide synthase (iNOS) and the anti-inflammatory marker arginase-1 (Arg-1). The proportion of iNOS-positive macrophages varied substantially among different cell-laden hydrogel formulations, ranging from 5% to 88% (Figure 5C). Similarly, the proportion of Arg-1-positive macrophages ranged from 4% to 92% across the AMGM hydrogel library (Figure 5D). The broad distribution of both markers suggests that differences in hydrogel composition and the associated matrix properties strongly influence MSC-mediated regulation of macrophage polarization.

To quantitatively evaluate MSC-mediated macrophage polarization across the AMGM hydrogel library, a standardized macrophage polarization index was adopted for subsequent machine-learning analysis. Consistent with a previously reported strategy, the index was calculated as log_2_[(M1/M2 ratio)×(M1 number+M2 number)], thereby integrating both the relative proportion and the absolute abundance of polarized macrophages into a single quantitative output [39].

The macrophage polarization index ranged from -3.95 to 4.02, with negative values indicating M2-dominant polarization and positive values indicating M1-dominant polarization. The broad distribution of index values across the hydrogel library suggests substantial differences in MSC-mediated regulation of macrophage polarization among the various AMGM formulations (Figure 5E). This quantitative index was subsequently used as the biological output parameter for machine-learning analysis of the relationships between matrix properties and MSC-mediated macrophage responses.

### 2.3 Virtual Database Construction and Screening of AMGM Hydrogels for MSC-Mediated Macrophage Polarization

To identify AMGM formulations that maximize MSC-mediated immunomodulatory capacity, regression models were established to correlate hydrogel composition with the macrophage polarization index, and the best-performing model was subsequently selected for virtual database construction and formulation screening (Figure 6A) [39]. Six algorithms were evaluated, including decision tree regression (DT), Gaussian regression (GR), support vector machine regression (SVM), least squares boosting regression (LSBoost), multilayer perceptron (MLP), and generalized additive model regression (GAM) [40–43].

**Figure 6.**
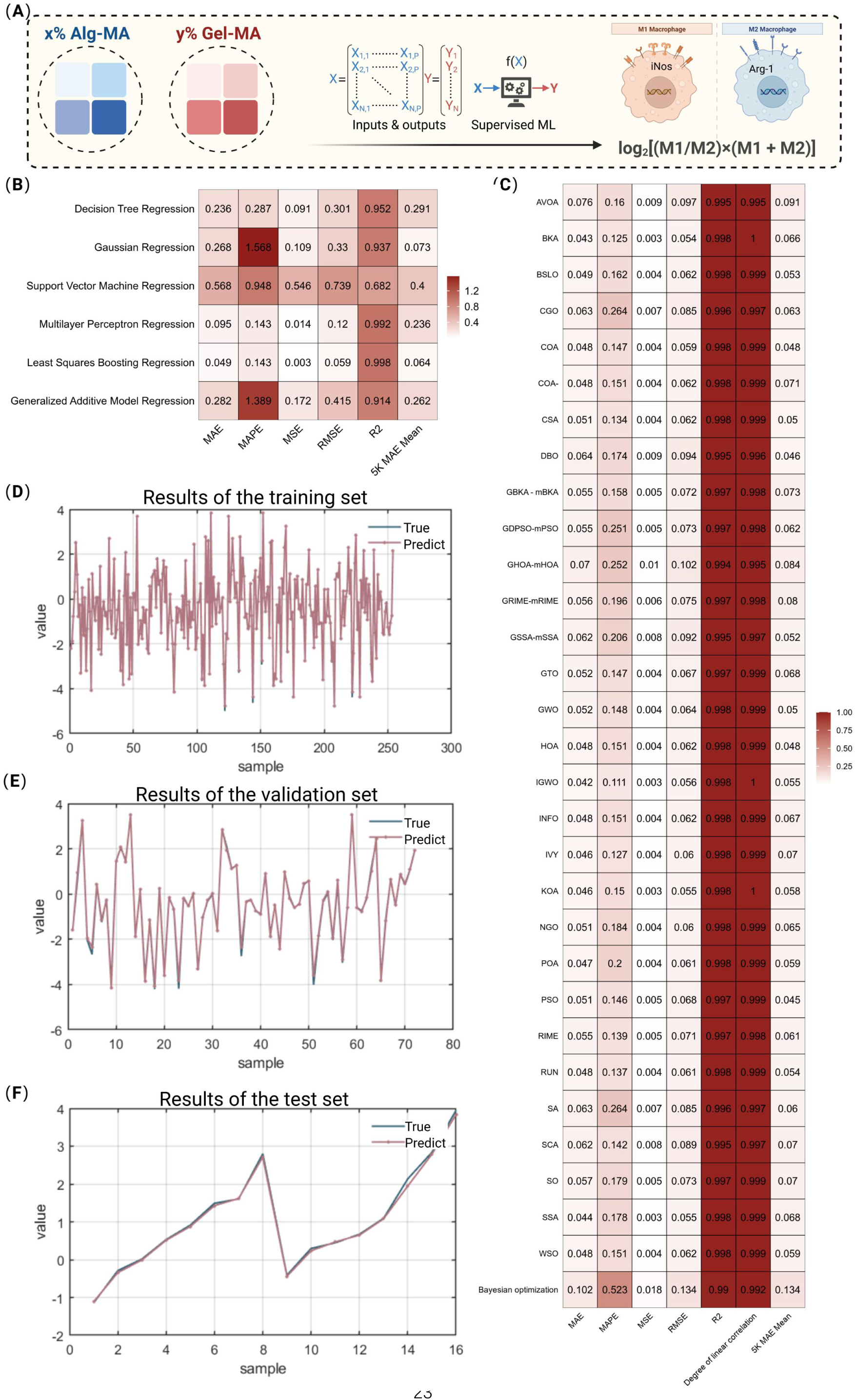
Machine-learning-based optimization and virtual screening of the AMGM hydrogel database. (A) Schematic illustration of the machine-learning workflow used to identify optimal AMGM hydrogel formulations for enhancing MSC-mediated macrophage polarization. (B) Performance comparison of six baseline machine-learning models and corresponding heatmap visualization of evaluation metrics. (C) Performance comparison of bayesian optimization together with 30 intelligent optimization algorithms applied to LSBoost and corresponding heatmap visualization of evaluation metrics. (D-F) Predictive performance of the PSO-optimized LSBoost model. Regression analyses comparing predicted and experimental values in the (D) training set, (E) validation set, and (F) test set.

Among these models, all except SVM achieved overall prediction accuracies above 90%. LSBoost showed the best predictive performance, with R² = 0.998 and a 5K MAE mean = 0.064 (Figure 6B). Accordingly, LSBoost was selected as the baseline model for subsequent optimization. Its superior performance is likely attributable to the ability of LSBoost to capture complex nonlinear relationships and improve robustness to dataset noise through residual-based optimization, making it well suited for modeling composition–function relationships within the AMGM hydrogel system [44].

We next optimized the LSBoost hyperparameters using Bayesian optimization together with 30 intelligent optimization algorithms. Among these approaches, particle swarm optimization (PSO) produced the best overall performance, yielding R² = 0.999 and a 5K MAE mean = 0.045 (Figure 6C, Supplementary Figure 4). The PSO-optimized LSBoost model was therefore selected as the final predictive model. The superior performance of PSO is likely attributable to its global search capability, which enables comprehensive exploration of the multidimensional parameter space and reduces the risk of convergence to local optima [45].

To further assess the robustness of the machine-learning framework and minimize the potential influence of replicate-level data leakage, an additional formulation-level analysis was performed. Specifically, technical replicates from the same hydrogel formulation were averaged, and each unique AMGM formulation was treated as an independent sample. Using this reformulated dataset comprising 54 unique hydrogel formulations, model training and validation were repeated following the same workflow. The optimized model retained strong predictive performance, yielding R² = 0.985 (Supplementary Figure 5), indicating that the primary predictive relationships were preserved even when replicate measurements were excluded from model construction.

To evaluate predictive performance, we compared predicted and experimental values in the training, validation, and test datasets. In all three datasets, the predicted values closely matched the experimental results, confirming the accuracy and stability of the optimized model (Figure 6D-F). Based on this model, we constructed a virtual database of AMGM formulations and screened candidate hydrogels with enhanced MSC immunomodulatory capacity. Together, these results establish a quantitative framework for predicting the immunomodulatory performance of binary AMGM MSC-laden hydrogels and provide a practical computational strategy for their screening and optimization.

### 2.4 Wet-Lab Validation of the Optimal Formulation for MSC-Mediated Macrophage Polarization Using the AMGM Hydrogel Database

To experimentally validate the predictions generated by the PSO-optimized LSBoost model, we performed wet-lab validation of the candidate formulations identified through virtual screening. Specifically, a high-throughput virtual screening strategy was implemented in which the compositional interval of the virtual hydrogel library was reduced to half of that used in the original experimental matrix, thereby generating a formulation space with finer compositional resolution and increased diversity. A refined validation matrix was subsequently constructed by reducing the concentration interval of both components in the binary AMGM system to half of the original spacing, allowing systematic evaluation of the computational predictions.

The macrophage polarization scores predicted by the virtual screening matrix showed strong agreement with the experimentally measured values, spanning from -3.62 to 3.98 (Figure 7A). Notably, formulations with identical total polymer concentrations did not necessarily exhibit comparable predicted immunomodulatory performance. For example, although AM_1_GM_0.5_ and AM_1.25_GM_0.25_ possess the same total polymer concentration (1.5% w/v), the latter exhibited a substantially lower predicted score, suggesting that the superior performance of AM_1_GM_0.5_ cannot be explained solely by reduced polymer density or a less restrictive network microenvironment. Ultimately, the PSO-optimized LSBoost model identified AM_1_GM_0.5_ as the optimal formulation.

**Figure 7.**
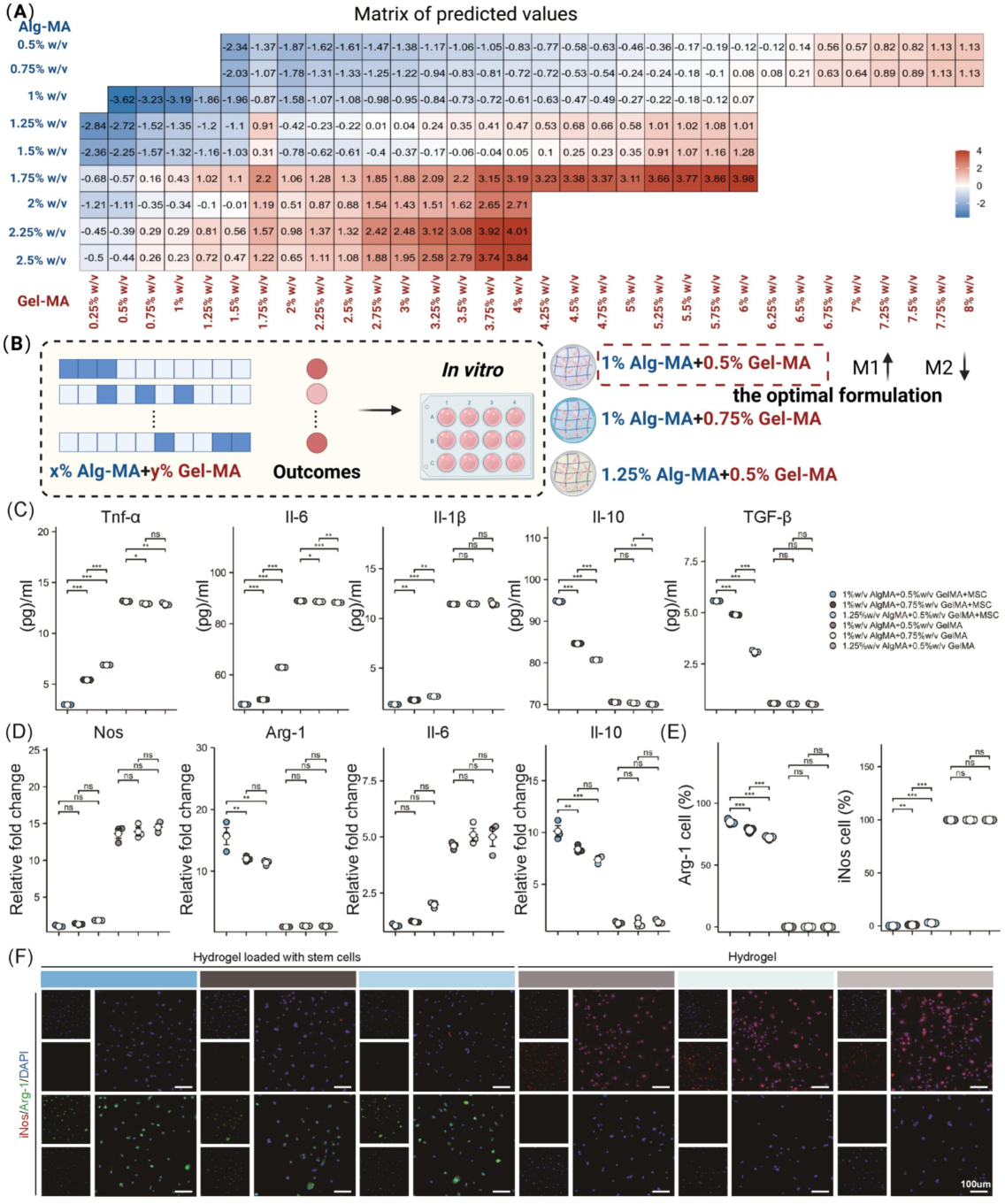
In vitro validation of the optimal AI-predicted formulation for MSC-mediated macrophage polarization. (A) Heatmap visualization of the prediction results from the PSO-optimized LSBoost model. (B) Schematic illustration of the in vitro validation strategy, including the predicted optimal formulation and neighboring validation formulations. (C) ELISA quantification of macrophage-secreted cytokines, including TNF-α, IL-6, IL-1β, IL-10, and TGF-β (n = 3). (D) qRT-PCR analysis of macrophage inflammation-related genes, including Nos2, Arg1, Il6, and Il10 (n = 3). (E) Quantitative analysis of immunofluorescence staining, showing the proportions of iNOS-positive and Arg-1-positive macrophages (n = 3). (F) Representative confocal immunofluorescence images showing the expression of the M1-associated marker iNOS and the M2-associated marker Arg-1 in macrophages (n = 3).

To verify the reliability of the model predictions, we performed in vitro validation of the predicted optimal formulation using a local compositional refinement strategy (Figure 7B). Starting from the predicted optimal formulation AM_1_GM_0.5_, neighboring candidate formulations were selected from the virtual database. Following a single-component variable adjustment principle, in which the concentration of one component in the binary AMGM system was adjusted while the other component was held constant, we selected AM_1.25_GM_0.5_ (increased AM, fixed GM = 0.5) and AM_1_GM_0.75_ (increased GM, fixed AM = 1) as neighboring validation formulations. After establishing this refined formulation array, immunomodulatory effects were evaluated by quantifying cytokine secretion from macrophages co-cultured with the different hydrogel formulations. ELISA analysis revealed significant differences in the secretion of pro-inflammatory cytokines (IL-1β, TNF-α, and IL-6) and anti-inflammatory cytokines (TGF-β and IL-10) among the experimental groups. Compared with the two neighboring formulations, the AM_1_GM_0.5_ group exhibited significantly lower levels of pro-inflammatory cytokines and significantly higher levels of anti-inflammatory cytokines (Figure 7C). In addition, all stem cell-laden hydrogel groups showed markedly different cytokine profiles compared with the corresponding cell-free material controls, confirming that the observed biological effects arose from the interaction between encapsulated MSCs and the hydrogel microenvironment rather than from the material composition alone.

To further validate these findings at the transcriptional level, quantitative real-time polymerase chain reaction (qRT-PCR) was performed to quantify the expression of representative pro-inflammatory genes (Nos2 and Il6) and anti-inflammatory genes (Arg1 and Il10). Significant differences in gene expression were observed between the MSC-laden hydrogel groups and the corresponding cell-free hydrogel controls, further demonstrating the immunomodulatory effects mediated by encapsulated MSCs. Among the three MSC-laden formulations, no significant differences were detected in the expression of the pro-inflammatory genes NOS2 and IL-6. In contrast, the expression levels of the anti-inflammatory genes Arg-1 and IL-10 were significantly higher in the AM_1_GM_0.5_ group than in the other two formulations, suggesting enhanced anti-inflammatory activity associated with the optimal formulation (Figure 7D).

Consistent with the qRT-PCR results, immunofluorescence analysis demonstrated superior immunomodulatory performance of the MSC-laden hydrogel groups compared with the corresponding cell-free hydrogel controls. Furthermore, although the proportion of iNOS-positive macrophages was comparable among the three MSC-laden formulations, the AM_1_GM_0.5_ group exhibited a significantly higher proportion of Arg-1-positive macrophages than the other two groups (Figure 7E, F). These findings were in good agreement with the transcriptional analyses and further supported the enhanced macrophage-regulating capacity of the AM_1_GM_0.5_ formulation.

To validate the practical utility and discriminatory power of the machine-learning-guided material optimization framework, we performed targeted experimental validation rather than re-screening the entire formulation space. Specifically, the model-predicted optimal formulation, AM_1_GM_0.5_, was compared with two neighboring formulations generated through single-component concentration adjustment, namely AM_1.25_GM_0.5_ and AM_1_GM_0.75_. This validation strategy was designed to assess whether the model could distinguish superior candidates within a narrow local composition space. In parallel, corresponding cell-free material controls were included to differentiate the effects of the optimized cell-laden microenvironment from those arising from material composition alone. Multidimensional in vitro analyses, including cytokine secretion, gene expression, and immunofluorescence characterization, consistently demonstrated that AM_1_GM_0.5_ exhibited the strongest capacity to promote anti-inflammatory macrophage polarization and suppress inflammatory responses. These findings support the predictive reliability of the machine-learning framework and its utility for data-driven biomaterial optimization (Figure 7).

To further investigate the in vivo immunomodulatory effects of MSC-laden hydrogels, we employed a rat subcutaneous implantation model, in which MSC-laden hydrogels of various formulations and their corresponding cell-free hydrogel controls were co-implanted into the dorsal subcutaneous region (Figure 8A). This parallel control design enabled comparison of the biological effects of the cell-laden constructs with those of the material alone [46]. Connective tissue thickness surrounding the implants was quantified at day 7, 14, and 21 after implantation as an indicator of the host inflammatory response (Figure 8B). The results showed that connective tissue thickness was significantly lower in the MSC-laden hydrogel groups than in the corresponding cell-free hydrogel controls, suggesting that the incorporation of MSCs attenuated the host inflammatory response following implantation. Histological analysis further revealed inflammatory cell infiltration surrounding the implants at day 7, which became more pronounced at day 14 and subsequently decreased by day 21 (Figure 8C-E). In addition, histological examination of major organs (Supplementary Figure 6A) and serum biochemical analyses (Supplementary Figure 6B-G) revealed no obvious abnormalities, indicating good in vivo biocompatibility and biosafety of the implanted hydrogels. These observations indicate a progressive resolution of the inflammatory response over time.

**Figure 8.**
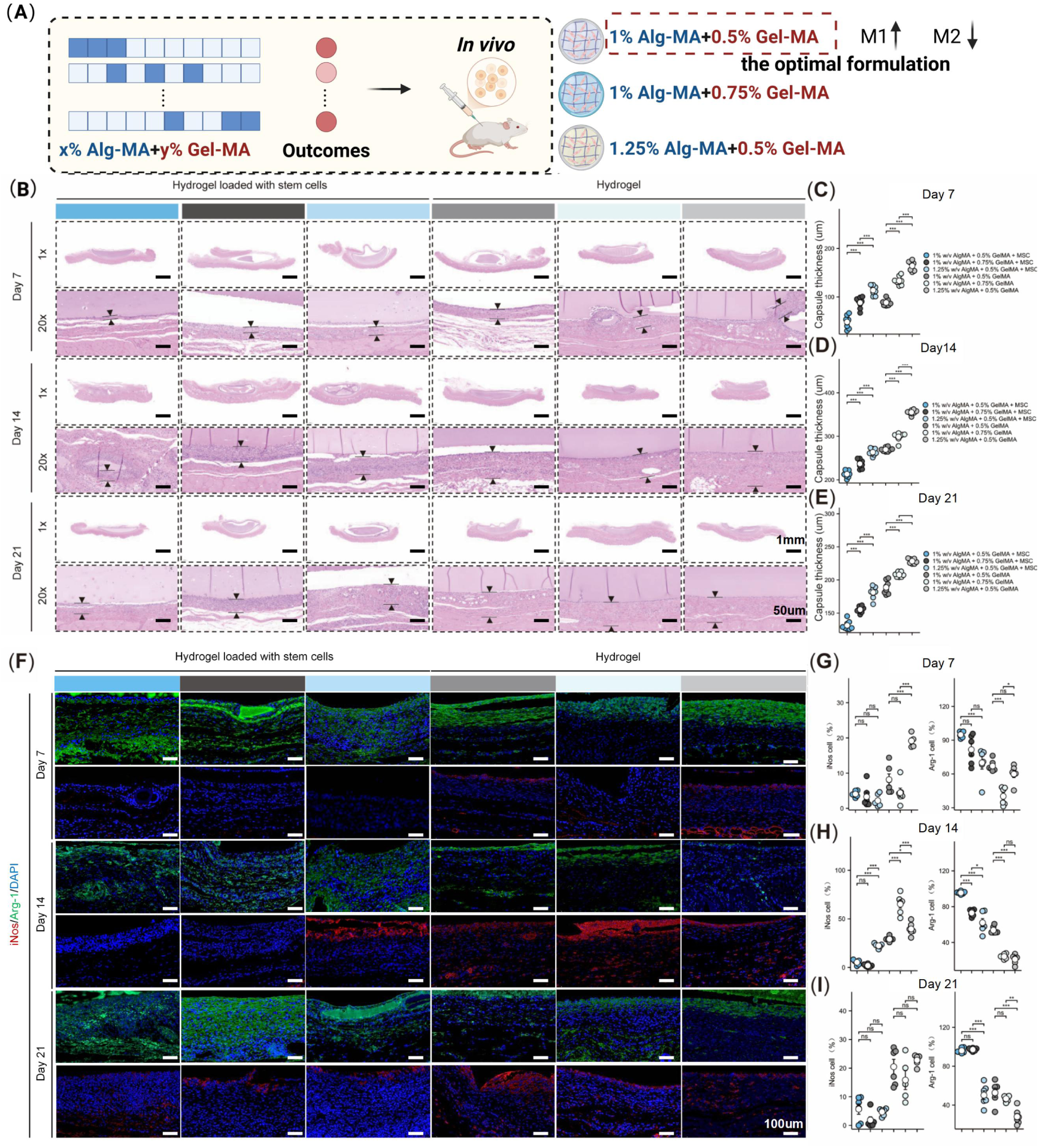
In vivo validation of the optimal AI-predicted formulation for MSC-mediated immunomodulation. (A) Schematic illustration of the in vivo experimental design, including animal grouping and implantation procedures. (B) Representative H&E-stained sections of peri-implant fibrous tissues collected at day 7, 14, and 21 after implantation. (C-E) Quantitative analysis of fibrous capsule thickness at (C) day 7, (D) day 14, and (E) day 21 post-implantation (n = 7). (F) Representative immunofluorescence images of peri-implant tissues stained for the M1-associated marker iNOS and the M2-associated marker Arg-1 at day 7, 14, and 21 after implantation. (G-I) Quantitative analysis of iNOS-positive and Arg-1-positive cells at (G) day 7, (H) day 14, and (I) day 21 post-implantation (n = 7).

To systematically evaluate the effects of MSC-laden hydrogels on the macrophage-associated immune microenvironment, macrophage infiltration and polarization were analyzed using established phenotypic markers. First, the proportion of infiltrating macrophages was quantified by flow cytometric analysis of CD11b and F4/80 expression (Supplementary Figure 7A). The results showed that the percentage of CD11b^+^F4/80^+^ macrophages was significantly lower in the MSC-laden hydrogel groups than in the corresponding cell-free hydrogel controls throughout the observation period (day 7, 14, and 21), suggesting reduced macrophage infiltration in the presence of MSCs (Supplementary Figure 7B-D).

Further analysis of tissue sections stained for the M1-associated marker iNOS and the M2-associated marker Arg-1 demonstrated that the stem cell-laden hydrogel groups exhibited a lower proportion of iNOS-positive macrophages and a higher proportion of Arg-1-positive macrophages than the corresponding cell-free hydrogel controls. Among the tested formulations, the stem cell-laden AM_1_GM_0.5_ hydrogel displayed the highest proportion of Arg-1-positive macrophages and the lowest proportion of iNOS-positive macrophages (Figure 8G-I).

Taken together, the fibrous capsule analysis and macrophage phenotypic characterization indicate that MSC-laden hydrogels modulate the local immune microenvironment following implantation. Among the tested formulations, AM_1_GM_0.5_ exhibited the most favorable macrophage polarization profile, consistent with its superior immunomodulatory performance observed in vitro.

### 2.5 SHAP-Based Feature Importance Ranking of Physicochemical Cues Governing MSC Immunomodulatory Capacity

To establish quantitative relationships between physicochemical properties of the hydrogel matrix and MSC-mediated immunomodulatory function, a machine-learning-based feature importance ranking framework was constructed using the physicochemical characteristics of the AMGM hydrogels and the corresponding macrophage polarization index data (Figure 9A; Supplementary Figure 8). First, recursive feature elimination with cross-validation (RFECV) was applied to evaluate the seven physicochemical features. The results indicated that pore size exhibited limited predictive value and was therefore excluded from subsequent analyses (Figure 9B). Subsequently, predictive models were trained using the remaining six physicochemical features together with the corresponding biological output data. Multiple regression algorithms, including DT, GR, SVM, LSBoost, MLP, and GAM, were evaluated. For each algorithm, the dataset was divided into training, validation, and test subsets for model optimization and performance assessment. Among all tested models, the multilayer perceptron (MLP) exhibited the best predictive performance, achieving R^2^ = 0.983 and a 5K MAE mean = 0.108 (Figure 9C). Accordingly, the MLP model was selected as the baseline model for subsequent analyses.

**Figure 9.**
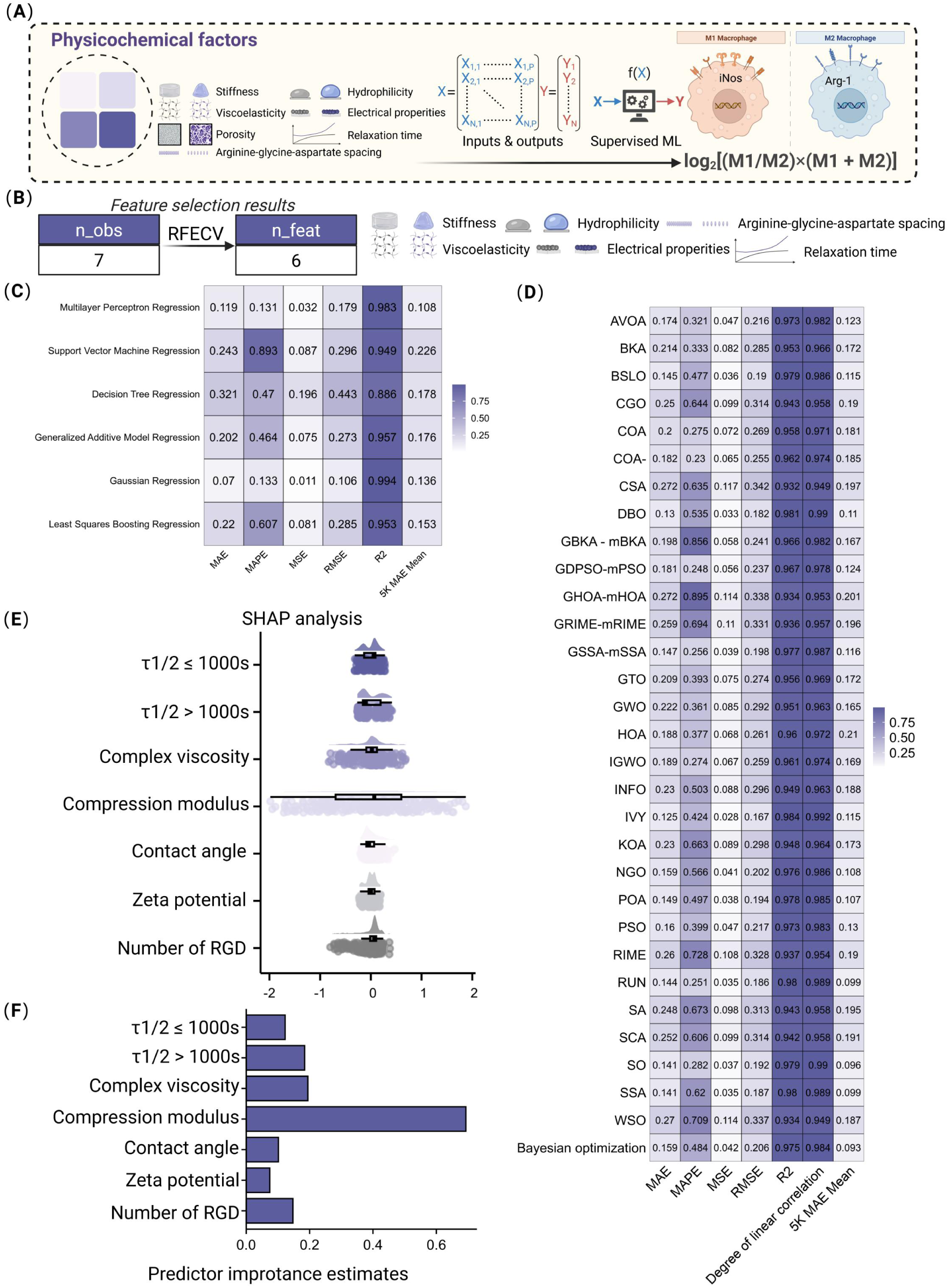
SHAP-based ranking of physicochemical feature importance in the AMGM hydrogel database. (A) Schematic illustration of the machine-learning workflow used to rank the relative importance of physicochemical features associated with MSC-mediated immunomodulatory function. (B) Feature selection using recursive feature elimination with cross-validation (RFECV). (C) Performance comparison of six baseline machine-learning models and corresponding heatmap visualization of evaluation metrics. (D) Performance comparison of 30 intelligent optimization algorithms applied to the MLP model and corresponding heatmap visualization of evaluation metrics. (E) SHAP summary plot showing the contributions of individual physicochemical features to model predictions across all samples. Positive SHAP values indicate positive contributions to the predicted output, whereas negative SHAP values indicate negative contributions. (F) Feature importance ranking based on the mean absolute SHAP values of all samples.

To further improve model robustness and generalization performance, the MLP model was subjected to systematic hyperparameter optimization after being selected as the baseline model. Hyperparameter tuning was performed using Bayesian optimization in combination with 30 intelligent optimization algorithms, and the performance of each approach was evaluated using 5-fold cross-validation [47]. The results showed that Bayesian optimization achieved the best overall performance, yielding R² = 0.984 and a 5K MAE mean = 0.093 (Figure 9D).

To visualize the contributions of physicochemical features to the predicted immunomodulatory performance of each hydrogel formulation, SHapley Additive exPlanations (SHAP) analysis was performed using the Bayesian-optimized MLP model. In the SHAP summary plot, positive SHAP values indicate positive contributions of a given feature to the predicted output, whereas negative SHAP values indicate negative contributions (Figure 9E). To quantify the relative importance of each feature, the mean absolute SHAP value was calculated. The results identified matrix stiffness as the most influential physicochemical parameter associated with MSC-mediated immunomodulatory function within the AMGM hydrogel system (Figure 9F).

This emphasis on matrix stiffness is consistent with a growing body of literature demonstrating that mechanical properties of the ECM can strongly influence cellular behavior and function. Mechanical cues have attracted increasing attention in cell biology and regenerative medicine, contributing to a paradigm shift in the design of biomaterials and cell-instructive microenvironments. Notably, the machine-learning predictions obtained in the present study are consistent with our previous findings showing that matrix stiffness can influence macrophage polarization and associated inflammatory responses [7]. Recent studies have further suggested that appropriate mechanical stimulation may mitigate MSC senescence and reduce inflammatory signaling [48]. Although these mechanisms were not directly investigated in the present work, they provide a potential biological context for understanding why matrix stiffness emerged as the highest-ranked predictive feature within the AMGM hydrogel system. Collectively, these findings support the interpretation that matrix stiffness represents a key predictive physicochemical parameter associated with MSC-mediated immunomodulatory function in this coupled material system.

The relative contributions of different microenvironmental cues to cellular behavior remain a topic of considerable debate. A wide range of matrix-associated properties have been shown to regulate cell fate and function. Among these, matrix stiffness is one of the most extensively studied parameters and has been widely implicated in controlling cell adhesion, cytoskeletal organization, force transmission, lineage specification, and paracrine activity, including the immunomodulatory functions of MSCs [7, 49]. In parallel, other mechanical descriptors, particularly those related to viscoelasticity, have also emerged as important regulators of cell behavior. A comprehensive review by Mooney and colleagues highlighted the association between ECM viscoelasticity and cellular functions, demonstrating that viscoelastic properties can spatially regulate cell morphology and temporally influence proliferation and differentiation [49, 50]. These effects arise because cells respond not only to the magnitude of matrix resistance but also to how that resistance evolves over time, as reflected by parameters such as stress relaxation, complex viscosity, and relaxation time. Beyond mechanical cues, other physicochemical properties, including adhesive ligand presentation and surface wettability, can also contribute to cell behavior. For example, Wei and colleagues demonstrated that RGD spacing strongly regulates stem cell adhesion and osteogenic differentiation, identifying ligand presentation as an important determinant of cell-matrix interactions [11]. Likewise, highly hydrophilic zwitterionic hydrogels have been shown to maintain the stemness of hematopoietic stem and progenitor cells in 3D culture, although the specific roles of hydrophilicity/hydrophobicity related properties in MSC-mediated immunomodulation remain less clearly defined [26].

Despite these diverse and often competing perspectives, previous studies have generally focused on individual parameters in isolation, and few have quantitatively compared the relative importance of multiple microenvironmental cues within a unified analytical framework. In the present AMGM hydrogel system, matrix stiffness emerged as the highest-ranked feature in the predictive model, whereas viscoelasticity and RGD spacing exhibited comparatively smaller contributions. However, this result should be interpreted within the context of the current material library rather than as a universal ranking of microenvironmental cues. In particular, the broader tunable range of stiffness in this system, together with the limited representation of relatively stiff yet fast-relaxing formulations, may have increased the apparent contribution of stiffness while reducing the ability to resolve the independent effects of viscoelasticity. Therefore, our findings support the importance of stiffness as the strongest predictive feature within the present material space, while also highlighting the need for more refined hydrogel designs to rigorously decouple stiffness, viscoelasticity, and other coupled matrix parameters (Supplementary Figure 9). Nevertheless, the present study establishes a useful framework for integrating machine learning with ECM formulation screening and provides a data-driven strategy for investigating multifactorial microenvironmental regulation.

Importantly, experimental validation within the present AMGM hydrogel system showed that formulations with higher matrix stiffness were associated with a progressive shift toward a pro-inflammatory macrophage polarization profile (Supplementary Figure 10A). At the transcriptional level, the expression of pro-inflammatory genes, including Nos2, Il6, and Tnfa, increased progressively, whereas the expression of anti-inflammatory genes, including Arg1, Il10, and Tgfb, decreased correspondingly (Supplementary Figure 10F). These trends were further supported at the protein level. Quantitative immunofluorescence analysis revealed a gradual increase in the proportion of iNOS-positive macrophages with increasing stiffness, accompanied by a corresponding decrease in the proportion of Arg-1-positive macrophages (Supplementary Figure 10B-D). Consistent with these findings, ELISA analysis demonstrated increased secretion of the pro-inflammatory cytokines IL-6 and TNF-α, together with reduced secretion of the anti-inflammatory cytokines IL-10 and TGF-β (Supplementary Figure 10E).

The strong agreement between the machine-learning predictions and the multidimensional experimental validation highlights the utility of this data-driven strategy for the design of MSC-laden hydrogels. Within the present AMGM hydrogel system, matrix stiffness emerged as the highest-ranked predictive feature associated with MSC-mediated immunomodulatory function. This finding provides a quantitative basis for prioritizing matrix stiffness during the design and optimization of MSC-laden hydrogels. More importantly, the present study establishes a generalizable analytical workflow in which a target biological function is first defined, translated into a quantitative endpoint, and subsequently linked to material composition and physicochemical properties through predictive modeling. This framework may facilitate the identification of high-priority design parameters before more focused mechanistic investigations, thereby improving the efficiency of biomaterial optimization.

## 3. Discussion

The biological performance of biomaterials used in tissue engineering is a critical determinant of regenerative outcomes, as local physicochemical cues within the extracellular microenvironment exert profound effects on the behavior of MSCs and immune cells [11]. However, because microenvironmental regulation is inherently complex, multidimensional, and nonlinear, conventional trial-and-error approaches often require extensive experimental effort and become increasingly inefficient as the number of variables increases. This challenge is particularly evident in the development of MSC-laden hydrogels, where both material and biological requirements must be considered simultaneously. On the one hand, the biomaterial must possess properties suitable for implantation, including biodegradability, processability, and low immunogenicity. On the other hand, the hydrogel microenvironment must support cell viability while maintaining the desired biological function.

A key translational contribution of the present study lies not only in the identification of an optimized hydrogel formulation but also in the establishment of a predictive framework linking material composition, physicochemical properties, and biological function. Through systematic in vitro and in vivo validation, we demonstrated that the machine-learning framework could reliably predict macrophage-associated immunomodulatory outcomes across the AMGM hydrogel library. Such a data-driven strategy may facilitate biomaterial screening, reduce reliance on extensive empirical testing and animal experiments, and accelerate the development of next-generation immunomodulatory biomaterials.

Forward design in the interdisciplinary fields of machine learning and polymer engineering increasingly relies on existing databases and virtual candidate libraries to facilitate target-oriented material screening [51, 52]. As research objectives become more specialized, publicly available datasets are often insufficient to support customized biomaterial design, motivating the development of application-specific databases tailored to particular functional objectives [53, 54]. While many existing studies focus primarily on predicting material properties, comparatively fewer investigations have attempted to predict biologically relevant cellular responses to biomaterials. This challenge is particularly pronounced for stem cell-laden hydrogels, where the generation of high-quality datasets requires cell encapsulation, long-term culture, and multidimensional biological characterization.

To improve predictive performance within the present study, several methodological strategies were implemented, including data augmentation to address limited sample size, RFECV-based feature selection to reduce feature redundancy, and intelligent optimization algorithms for model optimization. Together, these approaches enabled accurate prediction of macrophage-associated immunomodulatory outcomes within the AMGM hydrogel library. More importantly, the value of the present framework lies not only in predictive performance but also in its ability to connect material composition, physicochemical properties, and biological function within a unified analytical workflow. By enabling virtual screening prior to experimental validation, this strategy may improve the efficiency of biomaterial optimization, reduce experimental burden, and facilitate the rational design of MSC-laden hydrogels for regenerative medicine applications.

To improve the robustness and stability of the predictive framework, we implemented a systematic benchmarking strategy involving more than thirty intelligent optimization algorithms. Although the performance of individual metaheuristic algorithms can depend on the characteristics of a specific optimization problem, most algorithms showed satisfactory convergence behavior and stable predictive performance for the present multidimensional hydrogel dataset. This comparative analysis suggests that the identified material landscapes were not solely dependent on a single optimization strategy but rather reflected a relatively robust search process. Furthermore, SHAP analysis enabled quantitative evaluation of the relative contributions of individual physicochemical cues to MSC-mediated immunomodulatory function. These results suggest that conventional selection of microenvironmental factors based primarily on prior assumptions may overlook important interactions among coupled cues. In this regard, the three-layer materiomics framework proposed in the present study provides a useful strategy for integrating material composition, physicochemical properties, and biological function, thereby facilitating data-driven prioritization of candidate microenvironmental cues for subsequent mechanistic investigations.

While mechanotransduction has traditionally been studied using static matrices, native microenvironments are inherently dynamic and undergo continuous mechanical changes during tissue development, regeneration, and disease progression [55]. This limitation has stimulated growing interest in dynamic hydrogel systems with time-dependent properties [56–59]. In particular, photo-switchable matrices provide a powerful approach for actively modulating matrix stiffness and investigating cellular adaptation to evolving mechanical cues in real time [56–59]. Recent advances in photo-tunable hydrogels have further enabled the study of rapid stiffness switching and its effects on cellular mechanosensing [59]. Although the current AMGM platform represents a static hydrogel system, its measurable viscoelasticity may partially originate from the dynamic motion and physical entanglement of alginate-containing polymer chains, which can contribute to stress dissipation even in the absence of ionic calcium crosslinking. These observations further underscore the importance of incorporating matrix dynamics into the design of next-generation synthetic niches.

Moreover, our results suggest that matrix stiffness may serve as a useful design parameter for guiding subsequent optimization of MSC-laden hydrogels. Within the present material library, formulations with stiffness values close to 1 kPa exhibited favorable immunomodulatory performance, implying that future hydrogel design may benefit from prioritizing stiffness within this range rather than attempting to reproduce all physicochemical characteristics of the optimal formulation. Nevertheless, this interpretation should be considered within the constraints of the current material system. At the same time, we acknowledge several methodological limitations associated with physicochemical characterization. Conventional measurements of hydrophilicity/hydrophobicity are typically performed on 2D planar substrates and therefore may not fully capture the dynamic effects of hydrophilicity/hydrophobicity interactions experienced by encapsulated cells within 3D hydrogel networks. This limitation may partially explain the relatively low contribution of hydrophilicity/hydrophobicity-related parameters observed in the present data-driven model. Similarly, characterization of pore size using scanning electron microscopy of lyophilized hydrogels inevitably introduces structural deviations from the native swollen state. However, because the machine-learning framework relies primarily on relative differences among normalized features across formulations rather than absolute physical dimensions, these measurements remain suitable for comparative feature ranking within the current dataset.

Furthermore, although the human-to-mouse co-culture system used in this study has been extensively validated in several of our previous studies, we acknowledge that interspecies differences in cytokine signaling and receptor recognition efficiency may influence the fine-scale ranking of immunomodulatory performance.

Despite the predictive success of the present machine-learning framework, an important limitation arises from the inherent collinearity among material properties. In the binary AMGM hydrogel system, altering polymer composition inevitably induces simultaneous changes in multiple physicochemical cues, including stiffness, pore size, RGD ligand density, and viscoelasticity (Supplementary Figure S9). Consequently, the current experimental design does not allow complete physical decoupling of these parameters and therefore cannot establish their truly independent biological contributions. Accordingly, the term “decode” in this study should be understood as data-driven identification and prioritization of predictive microenvironmental descriptors within an experimentally defined material space, rather than as causal dissection of independently isolated physicochemical cues. In this context, the machine-learning model identifies matrix stiffness as the highest-ranked predictive feature and strongest correlative descriptor within the AMGM hydrogel system, but not as an independently validated causal regulator. More broadly, such feature collinearity represents a common challenge in polymer-based biomaterials. Under these conditions, machine learning can prioritize correlated features according to their predictive importance but cannot by itself assign fully independent biological effects. Future mechanistic studies will require orthogonal decoupling strategies to resolve the individual roles of microenvironmental cues with greater rigor.

The present findings should be interpreted within the design space defined by the AMGM material system investigated in this study. GelMA and AlgMA were selected because they are widely used biomaterials for stem cell delivery and possess favorable properties, including biocompatibility, biodegradability, ease of functionalization, and compatibility with remote photopolymerization. Moreover, as representative protein-and polysaccharide-based materials, respectively, they provide substantial physicochemical diversity and enable the construction of a hydrogel library with a relatively broad tunable range. Within this material space, stiffness spans a wider range than several other features and may therefore contribute more strongly to the current model output. Thus, rather than implying that stiffness is universally the dominant determinant across hydrogel systems, our findings suggest that matrix stiffness deserves particular consideration within the present material system when prioritizing future investigations. More importantly, the strength of the proposed framework lies in its ability to quantitatively rank the relative contributions of multiple coupled matrix features before more focused mechanistic studies. Because the entanglement of physical properties is an intrinsic characteristic of polymer networks, future materiomics studies will require multiscale decoupling strategies to rigorously isolate individual parameters and elucidate their independent biological mechanisms. Such experimental approaches, combined with data-driven feature prioritization, will be essential for deciphering the independent contributions of complex microenvironmental cues.

## 4. Conclusion

This study established a material-property-function correlation framework linking the physicochemical properties of biomaterials with MSC-mediated immunomodulatory function. Machine-learning approaches were employed to identify optimized formulations under coupled multifactorial conditions and to evaluate the relative contributions of physicochemical cues. Through systematic experimentation and model optimization, an optimized formulation for MSC-laden hydrogels was identified within the present material library. Importantly, the broader significance of this work lies not in the universal applicability of a specific hydrogel formulation, but rather in the demand-oriented methodological framework established herein. By defining a target biological function, selecting a quantitative functional endpoint, and constructing a material-property-function database, this workflow can, in principle, be extended to other biomaterial systems and customized design objectives. Future studies may focus on cross-platform validation in diverse material systems and the integration of deep learning approaches to establish more generalizable predictive frameworks based on the composition-property-function paradigm, thereby facilitating data-driven and precision-guided biomaterial design.

## 5. Methods

### 5.1 Cell isolation and Culture

#### 5.1.1 Isolation of mouse-derived macrophages

5-6 week old male BALB/c mice were purchased from Beijing Vital River Laboratory Animal Technology Co., Ltd. Mouse bone marrow-derived monocytes were isolated as previously described. Briefly, mouse bone marrow was flushed with RPMI 1640 medium and filtered through a 100 μm cell strainer (Jet, China). Bone marrow cells were resuspended at 6×10⁶ cells/mL in α-MEM (Hyclone, USA) supplemented with 10% fetal bovine serum (FBS, Gibco, USA), 1% penicillin-streptomycin (Hyclone, USA), and 30 ng/mL M-CSF (Peprotech, USA). Cells were cultured in a 100×20 mm treated petri dishes (Jet, China) for 5 days at 37 °C under a 5% CO₂ atmosphere, with fresh medium containing M-CSF added every 2 days. Subsequently, the purity of adherent macrophages was assessed by flow cytometry.

#### 5.1.2 Isolation of umbilical cord-derived mesenchymal stem cells

UCMSCs were isolated from Wharton’s jelly of the umbilical cord (No. 20240801, approved by the Ethics Committee of The Second Affiliated Hospital of Dalian Medical University, Dalian). The cells were cultured in growth medium consisting of α-MEM (Hyclone, USA), 10% FBS (Gibco, USA), and 1% penicillin/streptomycin (Hyclone, USA) at 37 °C in a humidified 5% CO₂ atmosphere, with medium replaced every 3 days. Subsequently, cells were passaged at 70–80% confluence and used at passage 5.

### 5.2 Preparation of Materials Omics

#### 5.2.1 Synthesis Steps of GelMA

GelMA was purchased from Huanova Biotech (Shenzhen, China). For preparation, 20 g of gelatin A (Sigma-Aldrich) was dissolved in 100 mL of phosphate buffer saline (PBS, pH 7.5) using a magnetic stirrer at 60 °C until fully homogenized. Then, 2 mL of methacrylic anhydride (MA, Sigma-Aldrich) was slowly added under vigorous stirring and mixed at 60 °C. The reaction was carried out for 3 h, after which the GelMA solution was transferred into a dialysis bag (MWCO 3500, with half of the bag volume left empty to facilitate water molecule exchange) and dialyzed against deionized water for 4 days (with water replaced three times daily), followed by lyophilization.

#### 5.2.2 Synthesis Steps of AlgMA

For AlgMA synthesis, 2 g of sodium alginate was dissolved in 100 mL of deionized water by stirring at 37 °C until completely dissolved, following a reported method.Then, 6 mL of methacrylic anhydride (MA) was then slowly added dropwise, with intermittent addition of 3 M NaOH to adjust the pH to 8.0. The reaction was conducted under an ice bath and protected from light for 24 h. The reaction solution was transferred into a dialysis bag (MWCO 3500, leaving half the volume empty for water exchange) and dialyzed against deionized water for 4 days (with the water replaced 3 times daily). After dialysis, the solution was centrifuged at 2000 g for 10 min to remove impurities, followed by vacuum filtration through 10 μm, 5 μm, 1.2 μm, 0.8 μm, 0.45 μm, and 0.22 μm hydrophilic mixed cellulose ester membranes sequentially. The purified solution was poured into trays, frozen at -80 °C overnight, and lyophilized in the dark for 4 days to obtain AlgMA.

#### 5.2.3 Composition ratio in materiomics

For hydrogel preparation, GelMA (0.5-10% w/v) and AlgMA (0.5-2.5% w/v) were dissolved in DPBS (Gibco, USA) according to the matrix design. A final concentration of 0.5% w/v of 2-hydroxy-1-[4-(2-hydroxyethoxy) phenyl]-2-methyl-1-propanone (Irgacure 2959, Sigma-Aldrich, UK) was added, followed by crosslinking under ultraviolet light (360 nm, 5 W/cm²) for 60 seconds.

### 5.3 Characterization of physicochemical properties

#### 5.3.1 Mechanical test

Before conducting stress-strain measurements, 800 μL of GelMA and AlgMA mixtures with different ratios were added into cylindrical plastic molds (diameter: 12 mm, height: 8 mm) in accordance with ISO 7743 standards, followed by UV curing for crosslinking. The compressive properties of GelMA and AlgMA hydrogels were tested using a mechanical tester (E43, MTS Instruments, USA) under a humidity conditions of over 60%. The elastic modulus was determined based on the initial slope of the linear region (2–8% strain) of the stress-strain curve.

#### 5.3.2 Rheological measurements

Rheological measurements were conducted using a hybrid rheometer (DHA, TA Instruments, USA). Samples were tested at 25 °C, with all measurements performed on a 20 mm flat steel plate geometry. To identify the linear viscoelastic region, amplitude strain sweep tests were initially carried out at a constant frequency (1 Hz) and variable shear strain (γ). Subsequently, oscillatory frequency sweep tests were performed at a constant strain of 1% to determine the storage modulus (G′) and loss modulus (G″) across a frequency range of 0.1-100 rad/s. For stress-relaxation analysis, each hydrogel sample was subjected to a 10% constant strain over a time period of 0-1000 s. The complex viscosity (η∗) was calculated using the formula:

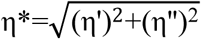

where η′ is the dynamic viscosity and η′′ is the imaginary viscosity, specifically defined as: Dynamic viscosity: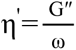, Imaginary viscosity: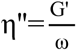. Here, G′ denotes the storage modulus, G″ denotes the loss modulus, and ω represents the angular frequency.

#### 5.3.3 Scanning Electron Microscopy

Hydrogels with different GelMA and AlgMA ratios were first frozen at -80 °C and then freeze-dried to remove moisture. To enhance their electrical conductivity, the hydrogels underwent gold sputtering. The pore sizes of hydrogels with varying compositions were observed using a scanning electron microscope (SEM) and quantified using ImageJ software.

#### 5.3.4 Characterization of zeta potentials

First, the mixed prepolymer solution was prepared in a PBS buffer system (pH 7.4) by adjusting the mass ratio of AlgMA and GelMA. Then, a 365 nm ultraviolet light source (with a radiation flux density of 5 mW/cm²) was used for the photo-induced cross-linking reaction. By precisely controlling the irradiation time at 60 seconds, the system underwent free radical polymerization to form a bulk hydrogel with a uniform 3D network. Subsequently, microsphere treatment was carried out using a JY92-IIDN ultrasonic cell disruptor. Under ice bath conditions, a pulse mode with 100 W power, consisting of 3 seconds of work followed by 3 seconds of interval, was set. The bulk hydrogel was then degraded into submicron-sized particles by the mechanical shear force induced by the cavitation effect. The zeta potentials of GelMA and AlgMA hydrogels were measured using a Zetasizer® Nano-S (NanoBrook 90Plus PALS) by laser Doppler electrophoresis at 25 °C after dispersing the hydrogels in HEPES buffer (5 mM, pH 7.0).

#### 5.3.5 Estimation of RGD density

Gelatin originates from the partial hydrolysis of collagen, a process that inherently preserves the native RGD cell adhesion sequences. Furthermore, the subsequent chemical modification with methacrylic anhydride to synthesize methacrylated gelatin does not compromise the structural integrity of these RGD motifs. Consequently, the total quantity of RGD ligands within the combinatorial AlgMA and GelMA hydrogels across varying precursor ratios was quantified based on a previously established mathematical model [60]. The specific calculation is formulated as follows.

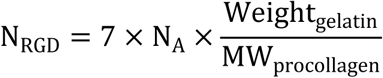

In this equation, *N_A_* represents Avogadros constant, Weight_gelatin_ denotes the mass of the gelatin component, and MW_procollagen_ indicates the molecular weight of procollagen, which is approximately 300 kDa [61].

#### 5.3.6 Hydrophilicity and hydrophobicity test

The hydrophilicity and hydrophobicity of AlgMA and GelMA hydrogels with different ratios were evaluated using the sessile drop method. A 10 μL droplet of deionized water was placed on the surface of the AlgMA and GelMA hydrogels, and the water contact angle was automatically calculated using a contact angle goniometer.

#### 5.3.7 Turbidity measurement

Hydrogel precursor solutions were prepared according to the corresponding AMGM formulations. Briefly, 200 μL of each precursor solution was transferred into a 96-well plate and allowed to stand at room temperature for 2 h. The turbidity of the precursor solutions was then evaluated using a microplate reader by measuring the absorbance at 600 nm. The absorbance values were subsequently visualized as heatmaps to illustrate the distribution of turbidity across different formulations.

### 5.4 Biocompatibility characterization of materials

#### 5.4.1 Cell viability staining

2×10⁵ MSCs were encapsulated in 200 μL of GelMA and AlgMA hydrogels and cultured for 24, 48, and 72 h, followed by viability assessment using a double staining method with Calcein-AM/EthD-1 (Invitrogen, USA). Live and dead MSCs were labeled with 2 mM calcein-AM and 4 mM PI, then incubated at 37 °C for 15 min. The cell live/dead staining was observed using a confocal laser scanning microscope (Olympus, FV3000, Japan).

#### 5.4.2 Cell proliferation

The proliferation of MSCs in 3D GelMA and AlgMA hydrogels was evaluated using a Cell Counting Kit-8 (CCK-8, Solarbio, China). A total of 2×10⁵ MSCs were seeded in 200 μL of GelMA hydrogels with varying stiffness in 24-well plates and cultured for 24, 48, and 72 hours. At the designated time points, 100 μL of the supernatant was transferred to 96-well plates, and 10 μL of CCK-8 solution was added to each well. Following incubation at 37 °C for 4 hours, the absorbance at a wavelength of 450 nm was measured using a microplate reader (Bio-Rad iMark, United States).

#### 5.4.3 Cell cytomorphology

2×10⁵ MSCs were encapsulated in 200 μL of GelMA hydrogels with different stiffnesses and cultured for 1, 2, and 3 days. For visualization of cell nuclei and F-actin, the samples were stained with 0.5 μg/mL 4′,6-diamidino-2-phenylindole (DAPI, Sigma-Aldrich, USA) for 5 minutes and with phalloidin (A12379, Thermo Fisher Scientific, USA, 1:400 dilution) for 1 hour at room temperature. The samples were washed twice with phosphate-buffered saline (PBS, Servicebio, China) and then observed using a confocal laser scanning microscope (FV3000, Olympus, Japan).

#### 5.4.4 BSA-FITC release analysis

To determine the release of proteins secreted by MSCs in GelMA and AlgMA hydrogels, 100 μg/mL FITC-labeled BSA (MW: 68 kDa, Solarbio, China) was mixed with 200 μL of GelMA hydrogel. The samples were then soaked in 1 mL PBS. At 0, 24, 48, and 72 hours, the supernatant was collected and transferred to a white, opaque 96-well plate. The BCA protein assay kit (Solarbio, China) was used to quantify the released proteins from GelMA hydrogels with different stiffness. The absorbance was measured at 562 nm using a microplate reader (Bio-Rad iMark, United States).

#### 5.4.5 Temporal framework for matrix construction and database generation

Generating the large-scale systematic dataset for the 162 combinatorial AMGM hydrogel matrices necessitated a rigorously phased experimental pipeline spanning several months. Initial material synthesis and lyophilization of the precursor polymers, specifically GelMA and AlgMA, required approximately one month of preparation. Subsequently, comprehensive profiling seven distinct physicochemical parameters across the entire 162 group library consumed two to three months of intensive analytical characterization. Finally, evaluating the downstream biological responses, particularly cellular biocompatibility and macrophage mediated immunomodulatory capacity, required an additional two to three months of robust in vitro experimentation.

### 5.5 Characterization of in vitro validation experiments

#### 5.5.1 Immunofluorescence staining

2×10⁵ MSCs were encapsulated in 200 μL of GelMA and AlgMA hydrogels, respectively, and co-cultured with 2×10⁵ macrophages in a Transwell co-culture system for 48 h. Macrophages were then fixed with 4% paraformaldehyde for 15 min at room temperature, washed three times with PBS, permeabilized with 0.1% Triton X-100 for 5 min, and blocked with 1% BSA for 30 min. Primary antibodies, including rabbit anti-Arginase-1 (Abcam, ab91279, Cambridge, MA; 1:200) and rabbit anti-iNOS (Abcam, ab178945, Cambridge, MA; 1:500) were incubated overnight at 4 °C. After washing with PBS, secondary antibodies Alexa Fluor 488-labeled goat anti-rabbit IgG (Invitrogen, Carlsbad, CA, 1:300) and Alexa Fluor 594-labeled goat anti-rabbit IgG (Invitrogen, Carlsbad, CA, 1:300) were applied in 1% BSA for 1 h at room temperature. Nuclei were counterstained with DAPI (Sigma-Aldrich, USA). Stained cells were visualized using a confocal laser scanning microscope (Olympus, FV3000, Japan). Images were analyzed using ImageJ software. Total cell counts were determined by nuclear staining, while M1 (iNOS+) and M2 (Arg-1+) macrophages were quantified based on fluorescence intensity thresholds.

#### 5.5.2 ELISA Detection of Cell Supernatants

The supernatant from the above co-culture system was collected to quantify the secretion of pro-inflammatory and anti-inflammatory cytokines. The levels of TNF-α, IL-6, IL-1β, IL-10 and TGF-β cytokines were quantitatively analyzed according to the ELISA kit instructions (Invitrogen, USA).

#### 5.5.3 Real-time quantitative reverse transcription

The concentration of extracted RNA was quantified using a NanoDrop 2000 spectrophotometer (Thermo Scientific), and cDNA was synthesized using the PrimeScript RT Reagent Kit with gDNA Eraser (RR047A, Takara, Japan). Primer sequences for the target genes Nos2, Arg1, Il6, Il10, and Actb are listed in Table 1. Quantitative PCR (qPCR) was performed with the following cycling conditions: initial denaturation at 95 °C for 30 s, followed by 40 cycles of 95 °C for 5 s, 56 °C for 30 s (annealing), and 72 °C for 20 s (elongation). Relative gene expression levels were calculated using the 2-ΔΔCt method, with Actb serving as the internal reference gene.

### 5.6 Characterization of in vivo validation experiments

#### 5.6.1 Subcutaneous implantation

Subcutaneous implantation animal experiments were approved by the Biomedical and Animal Ethics Committee of the Dalian University of Technology (DUTSBE250827-01). After anesthesia, 200 μL of AM_1_GM_0.5_, AM_1.25_GM_0.5_, and AM_1_GM_0.75_ solutions (with 2×10^5^ MSCs or without MSCs) were cross-linked with UV, and then randomly implanted on the left and right sides of the back of anesthetized male SD rats under sterile conditions. The operated rats were housed for 7 days, 14 days, and 21 days. The rats were euthanized by cervical dislocation. The implant and the adjacent area of full-thickness dermal tissue were excised together and immediately fixed in 4% paraformaldehyde for subsequent analysis.

#### 5.6.2 Histological staining

Retrieved subcutaneous explants were obtained and sectioned into 4 μm slices. Sections were dehydrated with a series of ethanol solutions of different concentrations. Afterward, the sections were treated with hematoxylin and eosin (HE).

#### 5.6.3 Histoimmunofluorescent staining

For immunofluorescent staining, CD11b+F4/80 antibodies (Servicebio, GB11058/GB11027, China; 1:200/1:5000), rabbit anti-iNOS antibody (Abcam, ab115819, Cambridge MA, USA; 1:500) and rabbit anti-Arginase-1 antibody (Abcam, ab91279, Cambridge MA, USA; 1:200) were incubated for overnights at 4 ℃, and the secondary antibodies, Alexa Fluor 488- and 594-labeled Goat anti-rabbit (Invitrogen, Carlsbad, CA) were incubated for 1 hour at room temperature. Finally, images were obtained with an upright optical microscope (NIKON ECLIPSE E100, Nikon).

### 5.7 Data Cleaning and Machine Learning Model Construction

#### 5.7.1 Data Processing

In the data preprocessing phase, we first applied the classic 3-sigma rule to identify outliers among the 162 original data points. Based on the theory of the normal distribution, this rule refines the dataset by calculating the mean and standard deviation, and eliminating data points that deviate more than three standard deviations from the mean. Subsequently, to establish a rigorous scientific model evaluation system, the integrated dataset was systematically ordered based on the concentration gradients of the two primary hydrogel components from low to high. Following this structured arrangement, the processed data were strictly partitioned into training, validation, and test sets at a ratio of 7:2:1. Given the inherent imbalance in data category distribution, the regression data within the training and validation sets were further clustered into six distinct categories.

To address the inherent challenge of data scarcity within specific minority categories, we implemented a sophisticated Generative Adversarial Network (GAN) architecture for strategic data augmentation. This model captured the underlying probability distribution of the original experimental data through a continuous adversarial game between a generator (G) and a discriminator (D). The generator was trained to transform random noise vectors from a normal distribution into synthetic samples that mimic the complex microenvironmental landscape, while the discriminator functioned as a binary classifier to distinguish between the original 162 biological data points and the generated counterparts [62].

The generative system was refined through the alternating optimization of two distinct loss functions via gradient descent. Specifically, the discriminator parameters were updated by minimizing the loss *L_D_*, defined as the negative expectation of the logarithmic probability for correctly identifying both real and synthetic samples. Simultaneously, the generator was optimized by minimizing its corresponding loss *L_G_* to effectively deceive the discriminator. This iterative training process continued until the performance of both networks reached an ideal equilibrium, allowing the synthetic data points to effectively cover the latent space of the AMGM hydrogel system. By functioning as an adversarial regularization strategy, this approach expanded the minority sample size to five times the original volume while strictly adhering to multidimensional physicochemical constraints and suppressing the risk of overfitting. To ensure that all physicochemical parameters possessed comparable scales and to eliminate the influence of disparate dimensions, a rigorous z-score standardization was performed on the augmented dataset prior to model training [63]. All physicochemical parameters were normalized.

#### 5.7.2 Feature Selection

REFCV serves as an advanced optimization of the Recursive Feature Elimination (RFE) method, integrating a cross-validation mechanism to establish a more rigorous feature selection framework [64]. This approach systematically iterates to remove features with lower contribution to the model, gradually reducing the dimensionality of the feature space to effectively avoid interference from redundant information and irrelevant variables in the data. In this process, cross-validation technology is used as a critical evaluation step. It conducts multiple rounds of training and validation on different feature subsets, using model performance metrics as quantitative criteria to precisely screen out the feature combination that maximizes the model’s generalization ability.

#### 5.7.3 Fundamental Machine Learning Regression Models

First, to measure the results of regression prediction, the following evaluation indicators are selected: MAE, MAPE, MSE, RMSE, and R² coefficient.

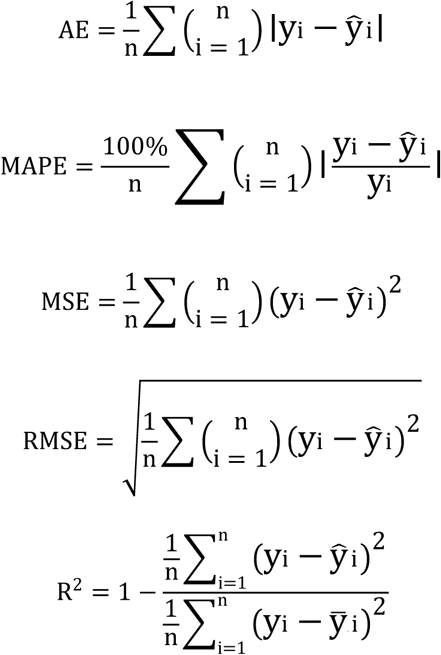

Let n be the sample size, yi be the true values, and ŷi be the predicted values. 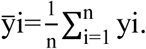.

LSBoost Regression is a regression algorithm based on the gradient boosting decision tree framework. It iteratively constructs a strong regression ensemble model by aggregating multiple weak regression base learners [43]. Each weak base learner is a decision tree that refines the model performance by fitting the residual errors of the current predictive model. LSBoost regression optimizes the model using the MSE loss function, which minimizes the squared error between the predicted and actual values.

GR, a non-parametric Bayesian regression method, utilizes Gaussian processes to probabilistically model the latent underlying function of input data [40]. It constructs a prior distribution through mean and covariance functions and updates it to a posterior distribution based on observed data for predictive inference. Defined as a probability distribution over an infinite-dimensional spaces of random functions, Gaussian processes characterize the uncertainty of function shapes with flexible priors. The squared exponential kernel (specified as ‘KernelFunction’,’squaredexponential’) is a typical covariance function widely used in this context. In MATLAB implementations, the fitrgp function is invoked to perform Bayesian updating of the prior distribution using input samples and corresponding outputs. The resulting posterior distribution is a Gaussian distribution, where the mean and covariance matrix provide the predictive values and uncertainty measures for new samples, respectively.

GAM regression, as a significant extension of nonlinear generalized linear models, achieves flexible modeling of the relationship between independent and dependent variables by decomposing the prediction function into a linear combination of smooth, nonlinear components [65]. This approach relaxes the restrictive assumptions of traditional linear models, leveraging basis function expansions to capture complex nonlinear mappings among variables. It retains the interpretability framework of generalized linear models while flexibly characterizing curvilinear trends and interaction effects using smoothing terms. In MATLAB implementations, the fitrgam function can be directly invoked to construct such models, which perform nonparametric estimation of variable effects using either automatically selected or user-specified basis functions.

DT, a non-parametric regression approach, recursively partitions the feature space into axis-aligned rectangular regions using decision tree algorithms, assigning the mean value of samples in each leaf node as the predicted value for continuous target variables [66]. Its tree-based structure inherently ensures high interpretability and excels at capturing nonlinear data relationships. However, this method is inherently prone to overfitting and instability. In practical applications, controlling tree complexity through pre-pruning or post-pruning techniques and integrating ensemble strategies such as Random Forest are essential to enhance generalization performance. Within the MATLAB environment, the fitrtree function can be directly invoked to construct a basic Decision Tree Regression model, enabling efficient modeling of complex data relationships through parameter optimization.

SVM is a nonlinear regression algorithm, which predicts continuous variables by constructing an optimal hyperplane and boundary band structure to map data into feature spaces [42]. Its core lies in maximizing the functional margin of samples closest to the hyperplane, ensuring that as many data points as possible fall within the boundary band. Leveraging the flexible mapping mechanism of kernel functions, this method efficiently handles nonlinear relationships in high-dimensional feature spaces, and it exhibits excellent generalization performance and strong robustness, making it especially suitable for small-sample and high-dimensional data scenarios. In practical applications, it is crucial to focus on the collaborative optimization of kernel function types and penalty term coefficients. The fitrsvm function in MATLAB enables convenient model construction, and combined with parameter tuning strategies such as grid search, it can significantly enhance the fitting accuracy and prediction reliability for complex data distributions.

As a typical feedforward neural network model, MLP is essentially a multi-layer nonlinear mapping system composed of several hidden layers and input/output layers [41]. Its basic computational unit is an artificial neuron that simulates the characteristics of biological neurons, enabling the layer-by-layer abstraction and high-order feature extraction of input features through weight connections and activation functions. In the MATLAB environment, an MLP model can be conveniently constructed by using the fitrnet function within the MATLAB Neural Network Toolbox.

#### 5.7.4 Machine Learning Intelligent Optimization Model Screening

In swarm intelligence optimization algorithms, remarkable innovations have been achieved in recent years, with a series of algorithms inspired by biological behaviors, physical phenomena, and mathematical principles. The Sparrow Search Algorithm (SSA, 2020) is dynamically modeled after the anti-predation behaviors of sparrows, achieving global optimization through role collaboration between scouts and followers in the population [67]. The Dung Beetle Optimizer (DBO, 2022) simulates the ball-rolling navigation, group interaction, and reproductive behaviors of dung beetles, constructing a robust solution space search mechanism based on celestial positioning [68]. The Sine Cosine Algorithm (SCA) designs search trajectories using the periodic oscillation characteristics of sine and cosine functions [69]. The Simulated Annealing (SA) algorithm balances exploration and exploitation via the Metropolis criterion, inspired by the energy decay principle of the physical annealing process [70]. The PSO algorithm achieves optimization iteration through information sharing among individual members in a bird swarm [71]. The Snake Optimization (SO) algorithm initiates a search process with a randomly initialized population [72]. The Pelican Optimization Algorithm (POA) enhances regional exploration and local fine-search capabilities by stage-wise simulation of surface reconnaissance and hunting behaviors [73]. The Grey Wolf Optimization (GWO) algorithm is based on the hierarchical structure and collaborative hunting patterns of wolf packs, while its improved version I-GWO effectively enhances population diversity and search balance by introducing a Dimension Learning Hunting strategy [74, 75].

Additionally, the African Vultures Optimization Algorithm (AVOA) [76], Chameleon Swarm Algorithm (CSA) [77], Gorilla Troops Optimizer (GTO) [78], Northern Goshawk Optimization (NGO) [79], and White Shark Optimizer (WSO) [80] draw inspiration from biological behaviors such as group foraging of birds, color adaptation of chameleons, social collaboration of gorillas, predation strategies of gannets, and sensory navigation of whale sharks, constructing optimization frameworks with distinct biological inspirations. Notably, the Chaos Game Optimization (CGO) integrates chaos theory and game theory [81]. The Information Weighted Mean Optimization (INFO) regulates search balance by weighting historical information [82]. The Coati Optimization Algorithm (COA) simulates hunting and escape behaviors [83]. The RIME algorithm constructs a search mechanism based on the process of ice growth over time [84]. The Kepler Optimization Algorithm (KOA) is inspired by the laws of planetary motion [85]. The RUNge-Kutta Optimizer (RUN) designs a metaphor-free search mechanism relying on numerical calculation methods [86]. The IVY algorithm models the growth and diffusion patterns of ivy plants [87]. The Hiking Optimization Algorithm (HOA) is modeled based on terrain elevation and walking functions [88]. The Black Kite Algorithm (BKA) combines Cauchy mutation and leader strategies [89]. The Crayfish Optimization Algorithm (COA) regulates three stages-avoid heat, competition, and foraging through temperature [90]. The Blood Sucking Leech Optimizer (BSLO) integrates five hunting strategies [91]. These algorithms all achieve the optimization process from random exploration to directional convergence in the solution space through the abstraction of natural phenomena, physical laws, or mathematical methods, providing diversified and efficient solution paradigms for complex scientific and engineering problems and demonstrating the interdisciplinary innovative and application potential in the field of swarm intelligence optimization.

#### 5.7.5 Assessment of Feature Independence and Interrelationships

To rigorously examine the dependencies and collinearity among the seven extracted physicochemical parameters, we performed a systematic pairwise correlation analysis. We utilized the raw experimental data points from all 54 hydrogel formulations to construct a correlation matrix. For each pair of variables, we computed the Pearson product-moment correlation coefficient to quantify the degree of linear association. The statistical significance of each correlation was assessed using a two-tailed Student’t test to identify statistically significant relationships. For the purposes of interpretation, we defined the degree of correlation based on the absolute value of the coefficient, where 0 to 0.3 indicated weak or no correlation, 0.3 to 0.5 suggested weak correlation, 0.5 to 0.8 represented moderate correlation, and 0.8 to 1 denoted strong correlation. These calculated coefficients along with their associated P-values are explicitly presented in the matrix plot to provide full transparency of the inherent couplings within the microenvironmental landscape.

### 5.8 Statistical analysis

Quantitative data were presented as mean values ± standard deviation (SD). Data were analyzed by the GraphPad Prism 5 and the statistical significance of the variance was evaluated by one-way analysis of variance (ANOVA) followed by Tukey’s multiple-comparison post hoc test. P < 0.05 was considered statistically significant.

## Ethics approval and consent to participate

The animal experiment was carried out in accordance with the Biomedical and Animal Ethics Committee of Dalian University of Technology (DUTSBE250827-01).

## Data availability

Data will be made available on reasonable request.

## Acknowledgements

Funding for this work was generously provided by the National Key Research and Development Program of China (No. 2022YFC2403002), the National Natural Science Foundation of China (No. 82501251, No. 52302344, No.52403152). Shenzhen Municipal Major Science and Technology Special Projects (KJZD20240903095722029), Liaoning Provincial Natural Science Foundation (Doctoral Research Startup Project 2025-BS-0995). Cross exploration of scientific research topics, medicine-engineering cross-joint fund (DUT25YG259).

## CRediT authorship contribution statement

Chenxi Pan: Conceptualization, Investigation, Methodology, Formal analysis, Data curation, Validation, Writing; Yi He: Investigation, Methodology, Data curation, Writing; Yingying Duan: Investigation, Methodology, Data curation; Kaiwen Chen: Conceptualization, Methodology, Formal analysis; Xiangying Wang: Resources, Data curation, Methodology, Formal analysis; Qifan Wang: Resources, Methodology; Yonggang Zhang: Supervision, Funding acquisition; Chuanfeng An: Supervision, Funding acquisition, Resources, Methodology, Validation; Huanan Wang: Conceptualization, Methodology, Investigation, Writing, Supervision, Project administration, Funding acquisition.

## Reference

[1] K. Le Blanc, D. Mougiakakos, Multipotent mesenchymal stromal cells and the innate immune system, Nature Reviews Immunology 12(5) (2012) 383–396.

[2] J. Doorn, G. Moll, K. Le Blanc, C. van Blitterswijk, J. de Boer, Therapeutic applications of mesenchymal stromal cells: paracrine effects and potential improvements, Tissue engineering. Part B, Reviews 18(2) (2012) 101–15.

[3] M. Najar, G. Raicevic, H. Fayyad-Kazan, D. Bron, M. Toungouz, L. Lagneaux, Mesenchymal stromal cells and immunomodulation: A gathering of regulatory immune cells, Cytotherapy 18(2) (2016) 160–71.

[4] D.M. Mosser, J.P. Edwards, Exploring the full spectrum of macrophage activation, Nature reviews. Immunology 8(12) (2008) 958–69.

[5] B.N. Brown, B.D. Ratner, S.B. Goodman, S. Amar, S.F. Badylak, Macrophage polarization: an opportunity for improved outcomes in biomaterials and regenerative medicine, Biomaterials 33(15) (2012) 3792–802.

[6] M.E. Wechsler, V.V. Rao, A.N. Borelli, K.S. Anseth, Engineering the MSC Secretome: A Hydrogel Focused Approach, Advanced healthcare materials 10(7) (2021) e2001948.

[7] Z. Zhuang, Y. Zhang, X. Yang, T. Yu, Y. Zhang, K. Sun, Y. Zhang, F. Cheng, L. Zhang, H. Wang, Matrix stiffness regulates the immunomodulatory effects of mesenchymal stem cells on macrophages via AP1/TSG-6 signaling pathways, Acta biomaterialia 149 (2022) 69–81.

[8] L. Cai, R.E. Dewi, A.B. Goldstone, J.E. Cohen, A.N. Steele, Y.J. Woo, S.C. Heilshorn, Regulating Stem Cell Secretome Using Injectable Hydrogels with In Situ Network Formation, Advanced healthcare materials 5(21) (2016) 2758-2764.

[9] X. Wang, S. Li, C. Yan, P. Liu, J. Ding, Fabrication of RGD micro/nanopattern and corresponding study of stem cell differentiation, Nano letters 15(3) (2015) 1457–67.

[10] S. Metwally, U. Stachewicz, Surface potential and charges impact on cell responses on biomaterials interfaces for medical applications, Materials science & engineering. C, Materials for biological applications 104 (2019) 109883.

[11] X. Liu, M. Zhang, P. Wang, K. Zheng, X. Wang, W. Xie, X. Pan, R. Shen, R. Liu, J. Ding, Q. Wei, Nanoscale distribution of bioactive ligands on biomaterials regulates cell mechanosensing through translocation of actin into the nucleus, Proceedings of the National Academy of Sciences of the United States of America 122(10) (2025) e2501264122.

[12] O. Chaudhuri, L. Gu, D. Klumpers, M. Darnell, S.A. Bencherif, J.C. Weaver, N. Huebsch, H.P. Lee, E. Lippens, G.N. Duda, D.J. Mooney, Hydrogels with tunable stress relaxation regulate stem cell fate and activity, Nature materials 15(3) (2016) 326–34.

[13] E.M. Sussman, M.C. Halpin, J. Muster, R.T. Moon, B.D. Ratner, Porous implants modulate healing and induce shifts in local macrophage polarization in the foreign body reaction, Annals of biomedical engineering 42(7) (2014) 1508–16.

[14] X. Han, X. Yang, R. Zhang, K. Chen, C. An, D. Li, S. Guo, F. Shao, C. Li, Y. Shen, Y. Zhang, Q. Ying, H. Wang, Hydrogels with hierarchical hydrogen bonds enable tunable stress relaxation to direct macrophage-driven immunoregulation, Acta biomaterialia 209 (2026) 391–407.

[15] N.R. Chan, B. Hwang, B.D. Ratner, J.D. Bryers, Monocytes contribute to a pro-healing response in 40 μm diameter uniform-pore, precision-templated scaffolds, Journal of tissue engineering and regenerative medicine 16(3) (2022) 297–310.

[16] B. Basu, N.H. Gowtham, Y. Xiao, S.R. Kalidindi, K.W. Leong, Biomaterialomics: Data science-driven pathways to develop fourth-generation biomaterials, Acta biomaterialia 143 (2022) 1–25.

[17] A. Suwardi, F. Wang, K. Xue, M.Y. Han, P. Teo, P. Wang, S. Wang, Y. Liu, E. Ye, Z. Li, X.J. Loh, Machine Learning-Driven Biomaterials Evolution, Advanced materials (Deerfield Beach, Fla.) 34(1) (2022) e2102703.

[18] Y. Zhang, Y. Fu, T. Sun, W. Li, L. Yu, J. Ding, Skin Relevant Biomaterials from Wound Healing, Medical Aesthetics, Flexible Electronics to Artificial Intelligence and Beyond, Advanced materials (Deerfield Beach, Fla.) (2025) e12919.

[19] Y. Zhou, X. Ping, Y. Guo, B.C. Heng, Y. Wang, Y. Meng, S. Jiang, Y. Wei, B. Lai, X. Zhang, X. Deng, Assessing Biomaterial-Induced Stem Cell Lineage Fate by Machine Learning-Based Artificial Intelligence, Advanced materials (Deerfield Beach, Fla.) 35(19) (2023) e2210637.

[20] W. Zhang, Y. Rao, S.H. Wong, Y. Wu, Y. Zhang, R. Yang, S.K. Tsui, D.F.E. Ker, C. Mao, J.E. Frith, Q. Cao, R.S. Tuan, D.M. Wang, Transcriptome-Optimized Hydrogel Design of a Stem Cell Niche for Enhanced Tendon Regeneration, Advanced materials (Deerfield Beach, Fla.) 37(2) (2025) e2313722.

[21] D. Fan, X. Chen, S. Wang, J. Zhan, Y. Chen, H. Zhou, D. Li, H. Tang, Q. He, T. Chen, Machine Learning-Assisted Prediction of Photothermal Metal-Phenolic Networks, Angewandte Chemie (International ed. in English) 64(13) (2025) e202423799.

[22] Y. Li, Z. Xu, J. Wang, X. Pei, J. Chen, Q. Wan, Alginate-based biomaterial-mediated regulation of macrophages in bone tissue engineering, International journal of biological macromolecules 230 (2023) 123246.

[23] B.J. Klotz, D. Gawlitta, A. Rosenberg, J. Malda, F.P.W. Melchels, Gelatin-Methacryloyl Hydrogels: Towards Biofabrication-Based Tissue Repair, Trends in biotechnology 34(5) (2016) 394–407.

[24] P. Zorlutuna, N. Annabi, G. Camci-Unal, M. Nikkhah, J.M. Cha, J.W. Nichol, A. Manbachi, H. Bae, S. Chen, A. Khademhosseini, Microfabricated biomaterials for engineering 3D tissues, Advanced materials (Deerfield Beach, Fla.) 24(14) (2012) 1782–804.

[25] N. Annabi, J.W. Nichol, X. Zhong, C. Ji, S. Koshy, A. Khademhosseini, F. Dehghani, Controlling the porosity and microarchitecture of hydrogels for tissue engineering, Tissue engineering. Part B, Reviews 16(4) (2010) 371–83.

[26] T. Bai, J. Li, A. Sinclair, S. Imren, F. Merriam, F. Sun, M.B. O’Kelly, C. Nourigat, P. Jain, J.J. Delrow, R.S. Basom, H.C. Hung, P. Zhang, B. Li, S. Heimfeld, S. Jiang, C. Delaney, Expansion of primitive human hematopoietic stem cells by culture in a zwitterionic hydrogel, Nature medicine 25(10) (2019) 1566–1575.

[27] H. Mohammadi, P.D. Arora, C.A. Simmons, P.A. Janmey, C.A. McCulloch, Inelastic behaviour of collagen networks in cell-matrix interactions and mechanosensation, Journal of the Royal Society, Interface 12(102) (2015) 20141074.

[28] S.C. Jasinoski, B.D. Reddy, Mechanics of cranial sutures during simulated cyclic loading, Journal of biomechanics 45(11) (2012) 2050–4.

[29] A.J. Engler, S. Sen, H.L. Sweeney, D.E. Discher, Matrix Elasticity Directs Stem Cell Lineage Specification, Cell 126(4) (2006) 677–689.

[30] R. Sridharan, D.J. Kelly, F.J. O’Brien, Substrate Stiffness Modulates the Crosstalk Between Mesenchymal Stem Cells and Macrophages, Journal of Biomechanical Engineering 143(3) (2020).

[31] Y. Ji, J. Li, Y. Wei, W. Gao, X. Fu, Y. Wang, Substrate stiffness affects the immunosuppressive and trophic function of hMSCs via modulating cytoskeletal polymerization and tension, Biomaterials Science 7(12) (2019) 5292–5300.

[32] O. Chaudhuri, L. Gu, M. Darnell, D. Klumpers, S.A. Bencherif, J.C. Weaver, N. Huebsch, D.J. Mooney, Substrate stress relaxation regulates cell spreading, Nature Communications 6(1) (2015) 6365.

[33] A.E.M. Beedle, S. Garcia-Manyes, The role of single-protein elasticity in mechanobiology, Nature Reviews Materials 8(1) (2023) 10–24.

[34] A.E.X. Brown, R.I. Litvinov, D.E. Discher, P.K. Purohit, J.W. Weisel, Multiscale Mechanics of Fibrin Polymer: Gel Stretching with Protein Unfolding and Loss of Water, Science 325(5941) (2009) 741-744.

[35] X. Zhao, Multi-scale multi-mechanism design of tough hydrogels: building dissipation into stretchy networks, Soft Matter 10(5) (2014) 672–687.

[36] E.E. Charrier, K. Pogoda, R.G. Wells, P.A. Janmey, Control of cell morphology and differentiation by substrates with independently tunable elasticity and viscous dissipation, Nature Communications 9(1) (2018) 449.

[37] L. Renesme, K.D. Cobey, M.M. Lalu, T. Bubela, R. Chinnadurai, J. De Vos, R. Dunbar, D. Fergusson, D. Freund, J. Galipeau, E. Horwitz, M. Lê, M. Matthay, D. Moher, J. Nolta, G. Parker, D.G. Phinney, M. Rao, J.E.J. Rasko, P.R.M. Rocco, F. Rossi, M.R. Myles, S. Viswanathan, B. Thébaud, Delphi-driven consensus definition for mesenchymal stromal cells and clinical reporting guidelines for mesenchymal stromal cell–based therapeutics, Cytotherapy 27(2) (2025) 146–168.

[38] T.A. Wynn, K.M. Vannella, Macrophages in Tissue Repair, Regeneration, and Fibrosis, Immunity 44(3) (2016) 450–462.

[39] H.M. Rostam, L.E. Fisher, A.L. Hook, L. Burroughs, J.C. Luckett, G.P. Figueredo, C. Mbadugha, A.C.K. Teo, A. Latif, L. Kämmerling, M. Day, K. Lawler, D. Barrett, S. Elsheikh, M. Ilyas, D.A. Winkler, M.R. Alexander, A.M. Ghaemmaghami, Immune-Instructive Polymers Control Macrophage Phenotype and Modulate the Foreign Body Response In Vivo, Matter 2(6) (2020) 1564–1581.

[40] M.W.J.I.j.o.n.s. Seeger, Gaussian Processes For Machine Learning, 14 2 (2004) 69–106.

[41] O.I. Abiodun, A. Jantan, A.E. Omolara, K.V. Dada, N.A. Mohamed, H. Arshad, State-of-the-art in artificial neural network applications: A survey, Heliyon 4(11) (2018) e00938.

[42] R.G. Brereton, G.R. Lloyd, Support Vector Machines for classification and regression, Analyst 135(2) (2010) 230–267.

[43] T. Hothorn, Boosting – An Unusual Yet Attractive Optimiser, Methods of Information in Medicine 53(06) (2014) 417–418.

[44] H.F. Jerome, Greedy function approximation: A gradient boosting machine, The Annals of Statistics 29(5) (2001) 1189–1232.

[45] M.S. Alajmi, A.M. Almeshal, Least Squares Boosting Ensemble and Quantum-Behaved Particle Swarm Optimization for Predicting the Surface Roughness in Face Milling Process of Aluminum Material, 11(5) (2021) 2126.

[46] M.D. Swartzlander, A.K. Blakney, L.D. Amer, K.D. Hankenson, T.R. Kyriakides, S.J. Bryant, Immunomodulation by mesenchymal stem cells combats the foreign body response to cell-laden synthetic hydrogels, Biomaterials 41 (2015) 79–88.

[47] R. Kohavi, A study of cross-validation and bootstrap for accuracy estimation and model selection, Proceedings of the 14th international joint conference on Artificial intelligence - Volume 2, Morgan Kaufmann Publishers Inc., Montreal, Quebec, Canada, 1995, pp. 1137–1143.

[48] X. Liu, Y. Ye, Z. Li, L. Liao, Q.J.N.C. Wei, Mechanical rejuvenation of senescent stem cells and aged bone via chromatin remodeling, 17 (2026).

[49] K. Ye, X. Wang, L. Cao, S. Li, Z. Li, L. Yu, J. Ding, Matrix Stiffness and Nanoscale Spatial Organization of Cell-Adhesive Ligands Direct Stem Cell Fate, Nano letters 15(7) (2015) 4720–9.

[50] O. Chaudhuri, J. Cooper-White, P.A. Janmey, D.J. Mooney, V.B. Shenoy, Effects of extracellular matrix viscoelasticity on cellular behaviour, Nature 584(7822) (2020) 535-546.

[51] S. Abdulghani, G.R. Mitchell, Biomaterials for In Situ Tissue Regeneration: A Review, 9(11) (2019) 750.

[52] H.I. Labouta, N. Asgarian, K. Rinker, D.T.J.A.N. Cramb, Meta-Analysis of Nanoparticle Cytotoxicity via Data-Mining the Literature, (2019).

[53] P. Rawat, R. Prabakaran, S. Kumar, M.M. Gromiha, AbsoluRATE: An in-silico method to predict the aggregation kinetics of native proteins, Biochimica et Biophysica Acta (BBA) - Proteins and Proteomics 1869(9) (2021) 140682.

[54] Z. Haider Jaffari, H. Jeong, J. Shin, J. Kwak, C. Son, Y.-G. Lee, S. Kim, K. Chon, K. Hwa Cho, Machine-learning-based prediction and optimization of emerging contaminants’ adsorption capacity on biochar materials, Chemical Engineering Journal 466 (2023) 143073.

[55] P. Roca-Cusachs, V. Conte, X. Trepat, Quantifying forces in cell biology, Nature cell biology 19(7) (2017) 742–751.

[56] X. Zhang, C. Dong, W. Huang, H. Wang, L. Wang, D. Ding, H. Zhou, J. Long, T. Wang, Z. Yang, Rational design of a photo-responsive UVR8-derived protein and a self-assembling peptide–protein conjugate for responsive hydrogel formation, Nanoscale 7(40) (2015) 16666–16670.

[57] X. Wu, W. Huang, W.-H. Wu, B. Xue, D. Xiang, Y. Li, M. Qin, F. Sun, W. Wang, W.-B. Zhang, Y. Cao, Reversible hydrogels with tunable mechanical properties for optically controlling cell migration, Nano Research 11(10) (2018) 5556–5565.

[58] L. Liu, J.A. Shadish, C.K. Arakawa, K. Shi, J. Davis, C.A. DeForest, Cyclic Stiffness Modulation of Cell-Laden Protein-Polymer Hydrogels in Response to User-Specified Stimuli including Light, Advanced biosystems 2(12) (2018).

[59] J. Yang, P. Wang, Y. Zhang, M. Zhang, Q. Sun, H. Chen, L. Dong, Z. Chu, B. Xue, W.D. Hoff, C. Zhao, W. Wang, Q. Wei, Y. Cao, Photo-tunable hydrogels reveal cellular sensing of rapid rigidity changes through the accumulation of mechanical signaling molecules, Cell Stem Cell 32(1) (2025) 121–136.e6.

[60] S. Jiang, C. Lyu, P. Zhao, W. Li, W. Kong, C. Huang, G.M. Genin, Y. Du, Cryoprotectant enables structural control of porous scaffolds for exploration of cellular mechano-responsiveness in 3D, Nature Communications 10(1) (2019) 3491.

[61] F.H. Silver, D.E. Birk, Molecular structure of collagen in solution: comparison of types I, II, III and V, International journal of biological macromolecules 6(3) (1984) 125–132.

[62] I. Goodfellow, J. Pouget-Abadie, M. Mirza, B. Xu, D. Warde-Farley, S. Ozair, A. Courville, Y.J.M.P. Bengio, Generative Adversarial Nets, (2014).

[63] G.W. Snecdecor, W.G. Cochran, Statistical Methods, Wiley 1991.

[64] I. Guyon, J. Weston, S. Barnhill, V. Vapnik, Gene Selection for Cancer Classification using Support Vector Machines, Machine Learning 46(1) (2002) 389–422.

[65] T. Hastie, R. Tibshirani, Generalized Additive Models: Some Applications, Journal of the American Statistical Association 82(398) (1987) 371–386.

[66] M. Czajkowski, M. Kretowski, The role of decision tree representation in regression problems – An evolutionary perspective, Applied Soft Computing 48 (2016) 458–475.

[67] J. Xue, B. Shen, A novel swarm intelligence optimization approach: sparrow search algorithm, Systems Science & Control Engineering 8(1) (2020) 22–34.

[68] J. Xue, B. Shen, Dung beetle optimizer: a new meta-heuristic algorithm for global optimization, The Journal of Supercomputing 79(7) (2023) 7305–7336.

[69] S. Mirjalili, SCA: A Sine Cosine Algorithm for solving optimization problems, Knowledge-Based Systems 96 (2016) 120–133.

[70] S. Kirkpatrick, C.D. Gelatt, M.P. Vecchi, Optimization by Simulated Annealing, Science 220(4598) (1983) 671-680.

[71] Z.H. Zhan, J. Zhang, Y. Li, H.S.H. Chung, Adaptive Particle Swarm Optimization, IEEE Transactions on Systems, Man, and Cybernetics, Part B (Cybernetics) 39(6) (2009) 1362–1381.

[72] F.A. Hashim, A.G. Hussien, Snake Optimizer: A novel meta-heuristic optimization algorithm, Knowledge-Based Systems 242 (2022) 108320.

[73] P. Trojovský, M. Dehghani, Pelican Optimization Algorithm: A Novel Nature-Inspired Algorithm for Engineering Applications, Sensors, 2022, p. 855.

[74] S. Mirjalili, S.M. Mirjalili, A. Lewis, Grey Wolf Optimizer, Advances in Engineering Software 69 (2014) 46–61.

[75] M.H. Nadimi-Shahraki, S. Taghian, S. Mirjalili, An improved grey wolf optimizer for solving engineering problems, Expert Systems with Applications 166 (2021) 113917.

[76] B. Abdollahzadeh, F.S. Gharehchopogh, S. Mirjalili, African vultures optimization algorithm: A new nature-inspired metaheuristic algorithm for global optimization problems, Computers & Industrial Engineering 158 (2021) 107408.

[77] M.S. Braik, Chameleon Swarm Algorithm: A bio-inspired optimizer for solving engineering design problems, Expert Systems with Applications 174 (2021) 114685.

[78] B. Abdollahzadeh, F.S. Gharehchopogh, S.J.I.J.o.I.S. Mirjalili, Artificial gorilla troops optimizer: A new nature-inspired metaheuristic algorithm for global optimization problems.

[79] M. Dehghani, S. Hubalovsky, P.J.I.A. Trojovsky, Northern Goshawk Optimization: A New Swarm-Based Algorithm for Solving Optimization Problems, 9 (2021) 162059–162080.

[80] M. Braik, A. Hammouri, J. Atwan, M.A. Al-Betar, M.A. Awadallah, White Shark Optimizer: A novel bio-inspired meta-heuristic algorithm for global optimization problems, Knowledge-Based Systems 243 (2022) 108457.

[81] S. Talatahari, M. Azizi, Chaos Game Optimization: a novel metaheuristic algorithm, 54(2 %J Artif. Intell. Rev.) (2021) 917–1004.

[82] I. Ahmadianfar, A.A. Heidari, S. Noshadian, H. Chen, A.H. Gandomi, INFO: An efficient optimization algorithm based on weighted mean of vectors, Expert Systems with Applications 195 (2022) 116516.

[83] M. Dehghani, Z. Montazeri, E. Trojovská, P. Trojovský, Coati Optimization Algorithm: A new bio-inspired metaheuristic algorithm for solving optimization problems, Knowledge-Based Systems 259 (2023) 110011.

[84] H. Su, D. Zhao, A.A. Heidari, L. Liu, X. Zhang, M. Mafarja, H. Chen, RIME: A physics-based optimization, Neurocomputing 532 (2023) 183–214.

[85] M. Abdel-Basset, R. Mohamed, S.A.A. Azeem, M. Jameel, M. Abouhawwash, Kepler optimization algorithm: A new metaheuristic algorithm inspired by Kepler’s laws of planetary motion, Knowledge-Based Systems 268 (2023) 110454.

[86] I. Ahmadianfar, A.A. Heidari, A.H. Gandomi, X. Chu, H. Chen, RUN beyond the metaphor: An efficient optimization algorithm based on Runge Kutta method, Expert Systems with Applications 181 (2021) 115079.

[87] M. Ghasemi, M. Zare, P. Trojovský, R.V. Rao, E. Trojovská, V. Kandasamy, Optimization based on the smart behavior of plants with its engineering applications: Ivy algorithm, Knowledge-Based Systems 295 (2024) 111850.

[88] S.O. Oladejo, S.O. Ekwe, S. Mirjalili, The Hiking Optimization Algorithm: A novel human-based metaheuristic approach, Knowledge-Based Systems 296 (2024) 111880.

[89] J. Wang, W.-c. Wang, X.-x. Hu, L. Qiu, H.-f. Zang, Black-winged kite algorithm: a nature-inspired meta-heuristic for solving benchmark functions and engineering problems, Artificial Intelligence Review 57(4) (2024) 98.

[90] H. Jia, H. Rao, C. Wen, S. Mirjalili, Crayfish optimization algorithm, Artificial Intelligence Review 56(2) (2023) 1919–1979.

[91] J. Bai, H. Nguyen-Xuan, E. Atroshchenko, G. Kosec, L. Wang, M. Abdel Wahab, Blood-sucking leech optimizer, Advances in Engineering Software 195 (2024) 103696.

